# Divergent Excitability of GABAergic Neurons Derived from Bipolar Disorder Patients Shapes Energy Shifts of Network Dynamics, possibly mimicking mania and depression

**DOI:** 10.1101/2025.08.28.672780

**Authors:** Ashwani Choudhary, Omveer Sharma, Ran Ben Ezer, Utkarsh Tripathi, Idan Rosh, Tchelet Stern, Jose Djamus Birch, Abraham Nunes, Gregory Falkovich, Martin Alda, Shani Stern

## Abstract

Bipolar disorder (BD) is characterized by fluctuating mood states, yet the cellular and circuit-level mechanisms distinguishing lithium-responsive (LR) from non-responsive (NR) patients remain elusive. We derived dentate gyrus granule neurons and GABAergic interneurons from induced pluripotent stem cells (iPSCs) of BD patients stratified by lithium response, and from healthy controls. Using patch-clamp electrophysiology, we assessed intrinsic excitability. We further developed a computational model simulating large-scale neuronal networks based on patient-derived electrophysiological properties and ion channel conductance distributions. Granule neurons from both LR and NR patients exhibited hyperexcitability compared to controls. However, GABAergic neurons showed a striking divergence: LR neurons were hyperexcitable, while NR neurons were hypoexcitable. Transcriptomic profiling revealed distinct molecular signatures between NR and LR neurons, including dysregulation of GABA receptor genes (*GABRR1*). Computational simulations over 10,000 iterations, mimicking long-term network activity, revealed that dentate granule neuron changes alone failed to recapitulate netwrok shifts between global hyperexcitability and global hypoexcitability. Networks incorporating only granule neuron phenotypes entered persistent hyper- or hypoactive states, depending on lithium response. Remarkably, when GABAergic neuron phenotypes were added to the model, both LR and NR networks exhibited spontaneous transitions between high and low activity states, that may be associated with the mood episodes that the patients exhibit. Control networks did not show such bistability. We, therefore, conclude that GABAergic neuronal excitability is a key determinant of lithium responsiveness in BD and critically shapes the emergence of state-shift dynamics in neural networks. These findings suggest that restoring excitatory/inhibitory (E/I) balance via targeted modulation of interneuron function may offer novel therapeutic avenues for BD.

## INTRODUCTION

Neural networks composed primarily of excitatory and inhibitory neurons regulate processing associated with simple and complex behaviors^1^. Excitatory synaptic inputs by glutamatergic neurons drive activity in a neural network, while inhibitory synaptic inputs modulate this activity to preserve stability^2^. Recently, E/I imbalance has been linked to various neuropsychiatric conditions such as BD, Autism spectrum disorders (ASD), and Schizophrenia (SZ)^3–6^. E/I imbalances can cause neuronal dynamics to tilt toward excessive excitation or inhibition, which can deeply impact behavior and mental state^7^. Studies on modulating excitation in the mouse prefrontal cortex, for example, have shown altered neuronal processing with increased gamma oscillations resulting in social behaviors similar to those seen in human mental disorders^8^. Interestingly, strengthening inhibition has the potential to restore normal behaviors^2,8^. Although validating these findings on humans has been difficult until the recent emergence of iPSC-based models^9–11^.

BD, with 1-2% of the global occurrence, is characterized by recurrent episodes of mania and depression^12,13^. BD is clinically classified as Type I, which has at least one manic episode that is often accompanied by depressive episodes; and Type II, which has recurrent depressive and hypomanic episodes without severe mania^13^. Pharmacologically, BD patients are classified as LRs (∼30%) or NRs (∼70%), with lithium being the first-line treatment^14,15^. BD is highly polygenic, with over 200 genetic associations that overlap with ASD and SZ, including variations associated with lithium responsiveness^16–20^.

iPSC-derived neurons from BD patients have revealed critical cellular and molecular phenotypes, including dysregulated gene expression in the Wnt signaling pathway, impaired synaptic function, and reduced neurite outgrowth^9,14,21,22^. A hallmark finding in BD iPSC-derived excitatory neurons is hyperexcitability, with studies demonstrating that lithium selectively normalizes this phenotype in hippocampal dentate gyrus (DG) neurons derived from LR patients but not NR patients^9,14,23,24^. For example, Mertens et al. first reported lithium’s ability to rescue hyperexcitability in iPSC-derived hippocampal neurons from LR patients, while additional work identified distinct electrophysiological signatures in LR versus NR BD neurons, offering potential predictors of lithium response^14,15,25^. Complementing these cellular insights, neuroimaging studies during manic episodes show heightened activity in limbic regions (amygdala, ventral striatum, anterior cingulate cortex) and overactive reward circuitry (orbitofrontal cortex, ventral striatum), correlating with elevated mood and impulsivity^26,27^. However, while it is compelling to associate mania with hyperactivity and depression with hypoactivity and lower energy states, hyperactivity has also been observed in the limbic regions (anterior cingulate cortex and amygdala) among depressed individuals in an fMRI study, especially in the ventromedial affective network that is central to negative mood regulation^28,29^.

Computational models of BD excitatory neurons have further corroborated these findings, simulating dynamic transitions between hyper- and hypoexcitable states that suggest the basis of clinical mood episodes^14,24^. These previous computational simulations^14,23,25^ performed single neurons’ responses, and have not dealt with the changes in the response of a large neuronal network due to the altered single neuronal responses. Additionally, the overwhelming focus on excitatory neurons has largely overlooked the role of inhibitory GABAergic neurons and E/I imbalance, despite compelling evidence of GABAergic dysfunction in BD^30,31^. Notably, postmortem studies have revealed dysregulation of GABA and associated proteins in the hippocampus and cortex of BD patients^32–36^. Additionally, alterations in GABA levels in plasma and cerebrospinal fluid from BD patients have been reported^37,38^. Studies have also shown a genetic basis for GABAergic dysfunction in BD, with reports of polymorphisms in GABA receptor genes (GABRA3 and GABRA5) associated with BD^39^.

To address this, we generated GABAergic-enriched neuronal cultures from iPSCs derived from control, BD-LR, and BD-NR subjects. This enabled us to perform a comprehensive analysis of the electrophysiological properties of BD patient-derived inhibitory neurons and their contributions to neuronal network activity. LR GABAergic neurons were found to be significantly more excitable than control GABAergic neurons. However, the excitability of NR GABAergic neurons was severely and significantly reduced. Further, leveraging RNA sequencing (RNA-seq), we identified differentially expressed genes (DEGs) among control, LR, and NR GABAergic neurons, providing insights into the dysregulated molecular pathways, including PI-3K signaling pathway, calcium signaling pathway, axon guidance, Wnt signaling, etc.

We performed computational modeling integrating these cellular findings at the single-cell level (based on our previous model^23,24^) and then at the network-level dynamics in BD, focusing on the dentate gyrus (DG). The single-cell models incorporated a random walk^40^ algorithm to simulate realistic physiological changes that occur in the neurons over time, which yield the same statistical properties of real-life neurons. When running the model over 10,000 iterations (simulating 10,000 neuronal time units), LR neurons exhibited hyperactive states, and NR neurons exhibited hypoactive states, but neither of them exhibited both.

We then developed a multicellular DG model using the NEURON^41^ platform based on a previously published model ^42,43^. The DG networks simulated complex connectivity between excitatory granule cells, GABAergic interneurons, and additional neurons in the DG, based on DG cyto-architecture. The developed approach enabled the simulation of neurophysiological properties over long periods, bypassing the need for impractical long-term recordings for hundreds of days. Thus, based on the electrophysiological recordings of excitatory and inhibitory neurons for ∼ 30 days, we were able to simulate ion-channel conductance sequences with matching statistical properties. These sequences were subsequently integrated into our computational simulations of multicellular networks, facilitating the study of long-term neural dynamics.

Our *in-silico* model recapitulated the dynamic network states characteristic of BD, possibly mirroring high energy and low energy clinical states. Interestingly, the DG network without incorporating BD-related changes in GABAergic neurons could not mirror both states. Rather, LR networks mirrored high energy-like states over prolonged periods, and NR networks mirrored low energy-like states over prolonged periods. Only after incorporating the physiological changes that we measured in BD patient-derived GABAergic neurons (different between LR and NR GABAergic neurons), both LR and NR networks started cycling between high and low activiy states that may be correlated with mania and depression states in the patients.

## RESULTS

### Differentiation of BD and control iPSCs into GABAergic neurons

We differentiated iPSC lines from six BD type I patients (3 LR and 3 NR BD patients) and three control individuals (Fig 1a). These iPSC lines were fully characterized^14^, and electrophysiological and transcriptional studies of excitatory hippocampal (DG granule neurons and CA3pyramidal neurons) and additionally motor neurons derived from these lines have been described previously^14,21,23^.

**Figure 1.**
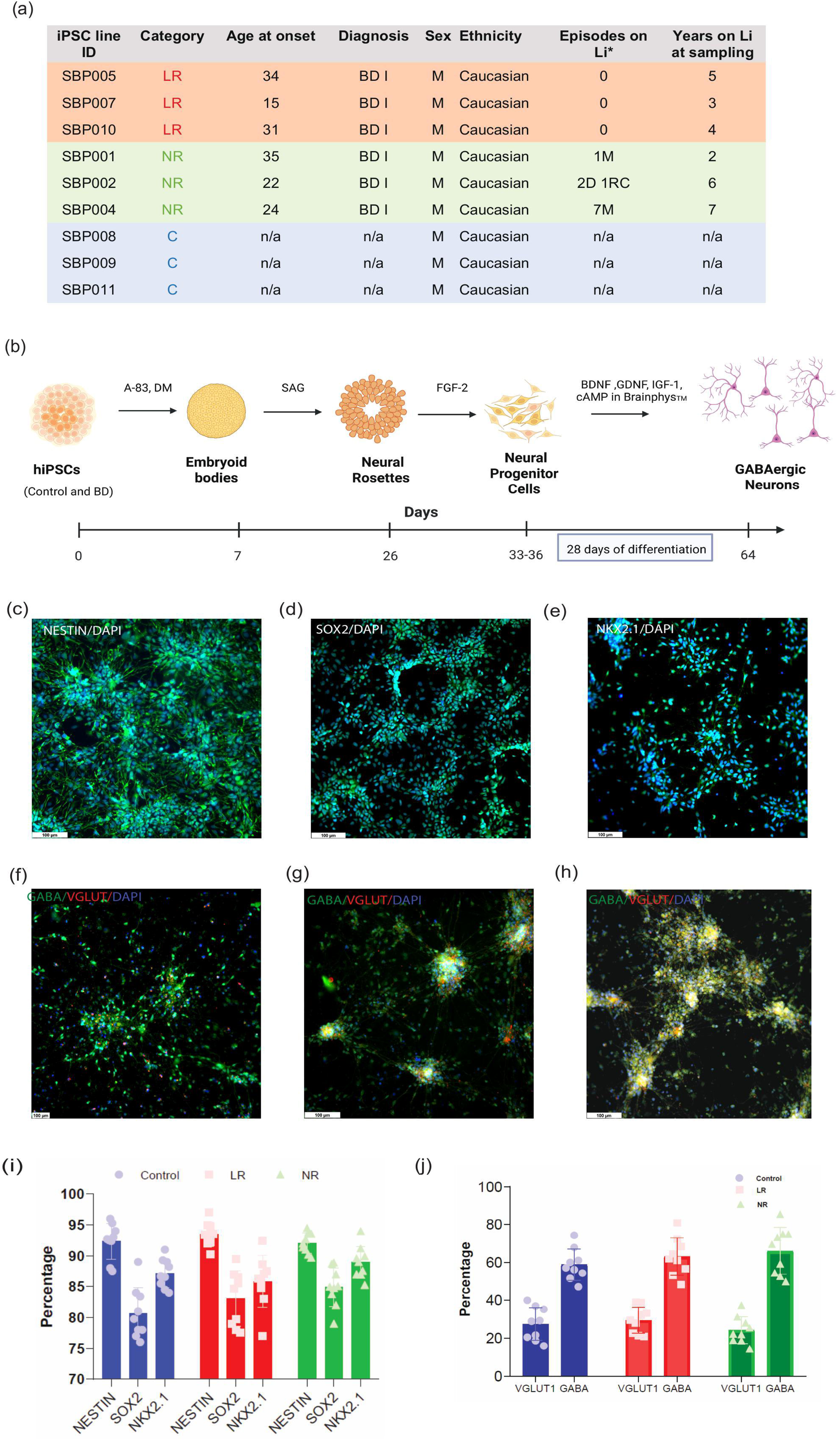
: Differentiation of BD and control iPSCs into GABAergic neurons. (a) Three iPSC lines from the control and three lines each from the LR and NR subtypes were used in this study. (b) Experimental design and detailed timeline for the differentiation of iPSC lines into GABAergic-enriched 2D *in vitro* neuronal culture. Immunocytochemistry of GABAergic NPCs with (c) NESTIN (green), (d) SOX2 (green), (e) NKX 2.1 (green), and DAPI (blue) for GABAergic NPCs (representative images). Immunocytochemistry of (f) control, (g) LR, and (h) NR stained with GABA (green), VGLUT-1 (Red), and DAPI (blue) (representative images). (i) Quantification of the immunostaining images for NESTIN, SOX-2, and NKX2.1 percentages in the three groups: control, LR, and NR. (j) Quantification of the immunostaining images for GABA, VGLUT percentages in the three groups: control, LR, and NR. *Data are presented as mean±SD*.

We successfully followed the recently published protocol for the differentiation of iPSCs into enriched cultures of GABAergic neurons^44^ with minor modifications (Fig 1b). During developmental stages, the excitatory neurons are mostly generated by dorsal forebrain progenitors, and the inhibitory GABAergic interneurons are generated by ventral forebrain progenitors^45^. Hence, iPSCs were first differentiated using small molecules like A-83-01 and dorsomorphin, promoting neuroectodermal commitment in embryoid bodies (EBs) via dual SMAD inhibition. Subsequently, Smoothened agonist (SAG) application activated *Sonic Hedgehog* signaling, enhancing ventral forebrain specification and driving rosette-stage cells toward GABAergic neuron precursors by upregulating ventral markers like NKX2.1 (Fig 1e). Finally, neurotrophic factors, along with specialized neuron medium^46^ (Brainphys^TM^), directed neural progenitor cells (NPCs) into GABAergic neurons. The detailed timeline and experimental design to obtain GABAergic neurons from the abovementioned iPSC lines are represented in Fig 1b. The NPCs were stained with relevant immunocytochemistry markers like NESTIN, SOX2, and NKX2.1 to ascertain their identity (Fig 1c-e). The percentage of NESTIN was 92±3%, 93±2%, and 92±2% in control, LR, and NR groups. Similarly, the percentage of SOX2 was 80±4%, 83±5%, and 85±3% in the three groups, respectively (Fig 1i). Additionally, the percentage of NKX2.1-expressing progenitor cells was 87±2%, 86±4%, and 89±3% in control, LR, and NR groups (Fig 1i). Further, to confirm the efficiency of the differentiation, we immuno-stained all the lines (including the control, LR, and NR groups) neurons respectively for GABA and VGLUT1 (Fig 1f-h). The percentage of GABA-expressing neurons was 58±8%, 63±10%, and 66±12% respectively, in control, LR, and NR. VGLUT1 was expressed in 28±9%, 30±7%, and 24±7% of the cells, respectively, in control, LR, and NR (Fig. 1j).

### Divergent Excitability Shifts in LR and NR GABAergic Neurons Relative to Controls

To investigate the functional properties of GABAergic neurons in BD, we differentiated iPSCs from 3 LR, 3 NR, and 3 control individuals into GABAergic neurons for 4-5 weeks. We performed whole-cell patch-clamp recordings to assess electrophysiological properties in LR, NR, and control neurons.

First, in voltage-clamp mode, we recorded sodium (Na+) and potassium (K+) currents. Normalized Naℒ currents were significantly larger in LR neurons compared to NR neurons but lower than in control neurons. Normalized fast and slow Kℒ currents showed no significant differences across groups. These findings suggest altered Naℒ channel function in BD GABAergic neurons, with LR neurons exhibiting intermediate currents between control and NR neurons (Fig 2a-c).

**Figure 2:**
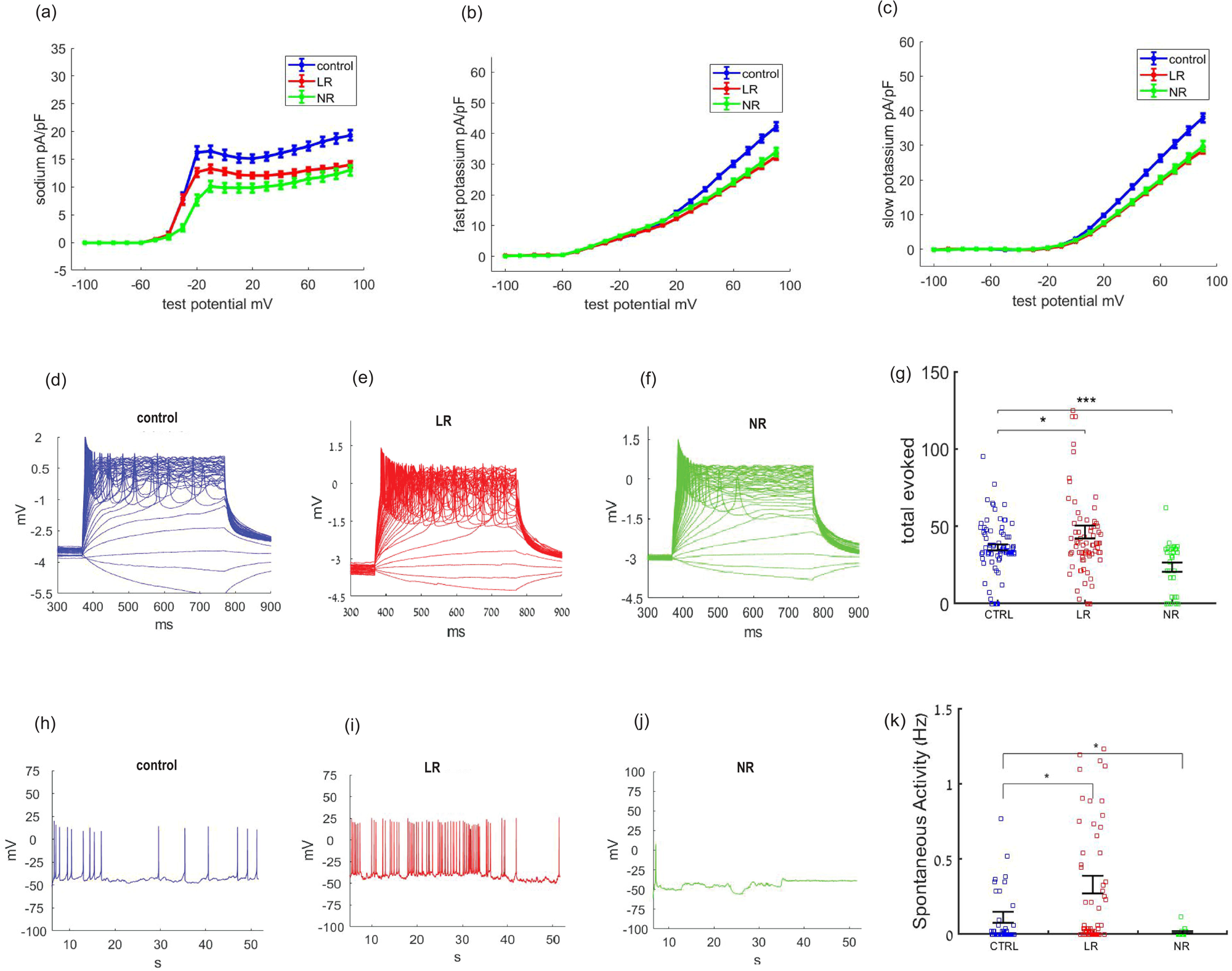
Divergent Excitability Shifts in LR and NR GABAergic Neurons Relative to Controls. Normalized currents by the capacitance (pA/pF) were plotted against different test potentials (- mV) in voltage-clamp mode. (a) The average Na+ currents for control (blue), LR (red), and NR (green) GABAergic neurons. Similarly, (b) the average slow K+ and (c) the average fast K+ currents are presented. Example traces for evoked action potentials of GABAergic neurons are shown in (d) control (blue), (e) LR (red), and (f) NR (green). (g) The average number of evoked potentials in the three groups. LR GABAergic neurons were hyperexcitable compared to controls, and NR GABAergic neurons were hypoexcitable compared to controls. Similarly, the example traces for spontaneous firing are shown in (h) control (blue), (i) LR (red), and (j) NR (green). (k) The average spontaneous firing rate. A significant decrease is observed in the firing rate of NR GABAergic neurons compared to controls, while a significant increase is observed in LR GABAergic neurons compared to controls. *Data are presented as mean±SEM*. \**p*<0.05, \*\**p*<0.01, \*\*\**p*<0.001.

We recorded evoked action potentials (APs) in current-clamp mode, holding neurons at -60 mV and injecting depolarizing currents (see *Methods*). LR neurons produced more evoked APs (46±4) compared to control (36±2) between ∼3-4 weeks of differentiation (p=0.03). NR neurons produced fewer APs compared to controls (24±3, p=8e-04) and were the least excitable of the three groups. It is interesting to note that, as DG granule neurons, both LR and NR neurons were hyperexcitable compared to control neurons^14^ and that LR CA3 pyramidal neurons were hyperexcitable compared to controls, but not NR CA3 pyramidal neurons. This further shows that BD neuronal alterations are cell-type specific. The severely reduced excitability of NR GABAergic neurons suggests a specific inhibitory deficit in NR BD patients (Fig 2d-f show representative examples and Fig. 2g presents the averages).

We next measured the spontaneous firing rate in current-clamp mode (see *Methods*) as an additional measure of neuronal excitability. LR GABAergic neurons exhibited a higher spontaneous firing rate (0.32±0.056 Hz) compared to controls (0.16±0.04 Hz) (p=0.025), while NR neurons exhibited a lower firing rate compared to controls (0.012±0.005 Hz) (p=0.029). The increased frequency of spontaneous firing rate in LR neurons further suggests intrinsic hyperexcitability, and the decreased frequency of NR neurons indicates their hypoexcitability compared to control GABAergic neurons (Fig 2h-j show representative examples and Fig. 2k presents the averages).

We further evaluated the effect of chronic lithium treatment on the excitability of BD GABAergic neurons (Supplementary Fig 1). Li+ treatment did not result in any significant change (p=0.3) in the excitability of LR neurons (40±9; evoked APs) compared to the LR neurons without treatment (36±2.3) (Supplementary Fig 1a-b show representative examples and Supplementary Fig 1c presents averages). Similarly, Li+ treatment did not affect the excitability of NR GABAergic neurons (20.2±2.8; evoked APs) in comparison to NR neurons without treatment (30±3) (p=0.23) (Supplementary Fig 1d-e show representative examples and Supplementary Fig 1f presents averages). Our previous reports^14^ show that chronic lithium treatment differentially affects DG neurons derived from patients who respond to lithium and does not affect DG neurons derived from non-responder patients. Here, we confirm that Li+ affects excitatory BD neuronal excitability but does not affect BD GABAergic neurons.

### Subtype-Specific Enrichment of Inhibitory Neuron Genes in LR and NR GABAergic Neurons Identified by Transcriptomic Profiling

RNA sequencing was next performed on control and patient-derived GABAergic neurons. Multiple genes were dysregulated between patients’ neurons and control neurons (the full DEG list is provided in Supplementary Table 1). GABAergic neuronal-related genes such as *VIP, FAT4*, *SLC6A20*, *GABRA1*, *GABRR1*, *GABBR2*, *CALCR*, and *KCNJ13* (∼678 DEGs; FDR<0.05) were found to be dysregulated in NR neurons compared to controls. Additionally, we found dysregulation of ventral marker genes (like *LHX5* and *DLX6*), and Wnt signaling pathway genes (*WNT5A*, *WNT8B*, *WNT7A*) in NR neurons (Fig 3a). When comparing NR and control GABAergic neurons seeking enriched KEGG pathway genes in NR neurons, notably neuroactive ligand-receptor interaction (63 genes, FDR=1.2E-6), calcium signaling (43 genes, FDR=8.3E-05), Wnt signaling (29 genes, FDR=0.003), and axon guidance (29, FDR=0.006) (Fig 3f) pathways were enriched (Fig 3b).

**Figure 3:**
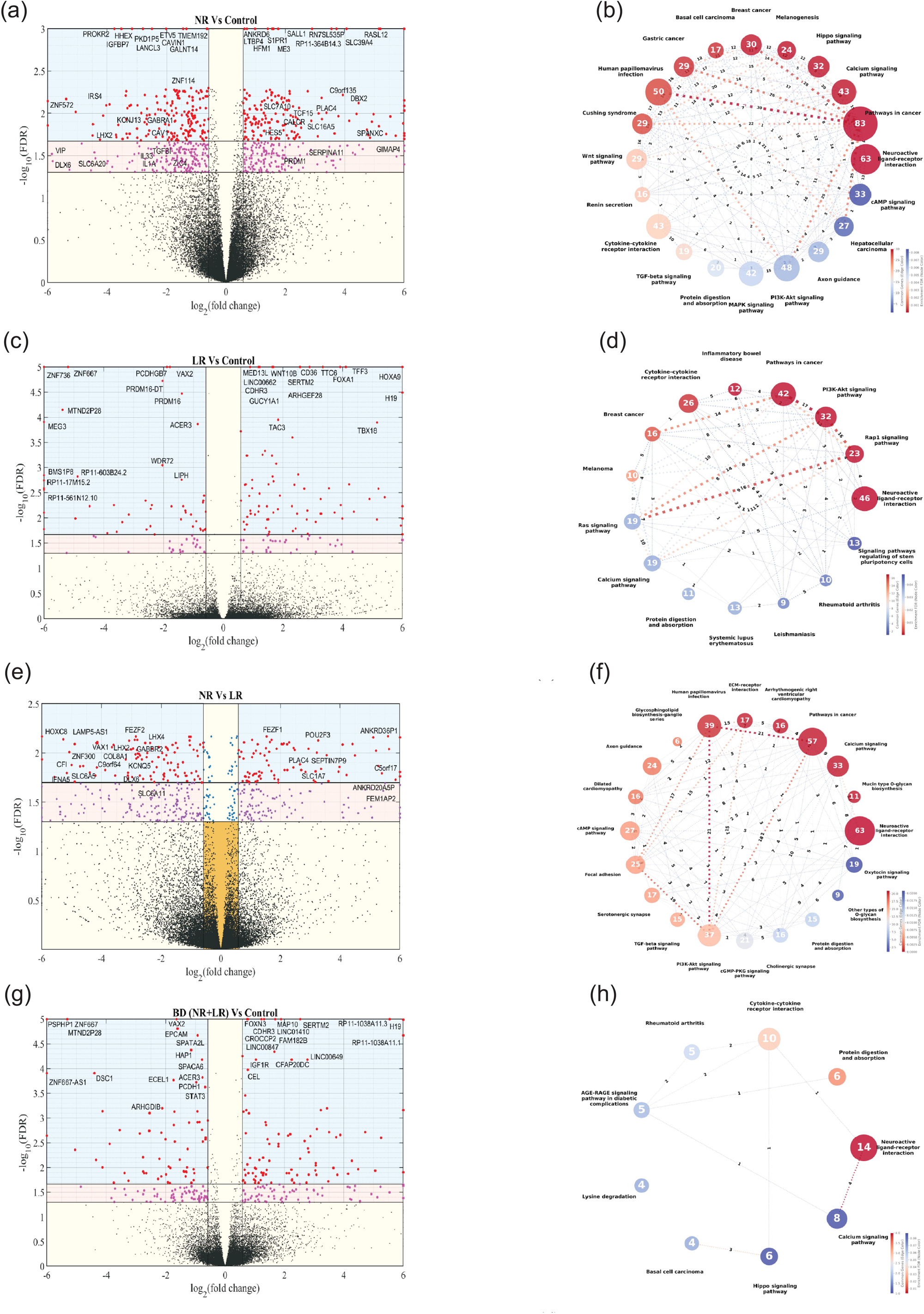
Subtype-Specific Enrichment of Inhibitory Neuron Genes in LR and NR GABAergic Neurons Identified by Transcriptomic Profiling. (a) A volcano plot for the differentially expressed genes between control and BD GABAergic neurons (including LR and NR subtypes) with |log2 fold changeL|L> 0.5 and FDR<0.05 (light purple dots). Genes with |log2 fold changeL|L> 0.5 and FDR<0.02 are represented in red dots. (b) The corresponding network analysis for the KEGG pathways of the differentially expressed genes *(p*-value <0.05) between control and BD neurons. (c) A volcano plot for the differentially expressed genes between control and LR GABAergic neurons; FDR and log2 fold change cut-off same as (a). (d) The corresponding network analysis for the KEGG pathways of the differentially expressed genes between control and LR neurons. (e) A volcano plot for the differentially expressed genes between control and NR GABAergic neurons; FDR and log2 fold change cut-off same as (a). (f) The corresponding network analysis for the KEGG pathways of the differentially expressed genes*(p*-value <0.05) between control and NR neurons. (g) A volcano plot for the differentially expressed genes between NR and LR GABAergic neurons. FDR and log2 fold change cutoff same as (a). (h) The corresponding network analysis for the KEGG pathways of the differentially expressed genes *(p*-value <0.05) between NR and LR neurons.

Approximately, 59 DEGs (FDR<0.05) including *GABRR1*, *GBE1*, *TRP6*, *IGFIR*, *ENO4*, *COL24A1*, *COL21A1*, and *PCDH8* were dysregulated in LR neurons compared to controls (Fig 3c). When comparing LR and control GABAergic neurons, LR neurons showed significant upregulation of KEGG pathways^47^ associated with synaptic function and plasticity, such as neuroactive ligand-receptor interaction (46 genes, FDR=7.7E-11), PI3K-Akt signaling (32 genes, FDR=2.3E-6), Rap1 signaling (23 genes, FDR=2.3E-6), and calcium signaling (19 genes, FDR=0.04) (Fig 3d). These pathways, which are essential for neuronal survival, synaptic remodeling, and excitability, may contribute to the hyperexcitability observed in LR GABAergic neurons^48–51^.

Specific comparisons between NR and LR GABAergic neurons provided more insights into the molecular variations underlying their contrasting neurophysiology. We found GABA receptor (like *GABBR2*), and *SLC6A11, SSTR2*, *LAMP5-AS1*, *VAX1*, *FAT4*, and *DLX6* dysregulated in the analysis of NR vs LR neurons (∼ 446 DEGs; FDR<0.05) (Fig 3e). Interestingly, ventrally expressed genes like *NKX2-2* and *NKX6-1* were also dysregulated between NR and LR

GABAergic neurons. NR neurons displayed stronger enrichment of PI3K-Akt signaling (37 genes, FDR=0.006), neuroactive ligand-receptor interaction (63 genes, FDR=7.2E-13), calcium signaling (33 genes, FDR=0.0003), and serotonergic synapse (17 genes, FDR=0.005). On the other hand, LR neurons showed a relative increase in cGMP-PKG signaling (21 genes, FDR=0.01) (Fig 3f) (Supplementary Fig 2).

*GABRA1*, *FAT3, VAX2, IGFIR*, *DSC1, BHLHE22, GIGYF1*, and *ZNF667* (total 296 DEGs; FDR<0.05) were found to be dysregulated in BD neurons when LR and NR subtypes were analyzed together as a group compared to controls (Fig 3g). KEGG pathway analysis of the BD group (LR and NR together) contrastingly showed lower enrichment of pathways such as Neuroactive ligand-receptor interaction, Calcium signaling pathways (Fig 3h).

Using Principal Component Analysis (PCA) based dimensionality reduction on a less stringent and broader set of DEGs (*p*<0.05, DEseq2; list of DEGs in Supplementary table 1), we observed distinct clustering patterns for the GABAergic neurons. LR GABAergic neurons clustered closer to control neurons in the first quadrant. In contrast, NR GABAergic neurons were distinctly positioned in the second quadrant, farther from the other two groups, indicating a unique transcriptomic signature in NR neurons that sets them apart from control and LR GABAergic neurons (Supplementary Fig 3).

### Long-term *in-silico* simulation of single-cell excitatory DG granule neurons incompletely captures the network dynamics associated with BD subtypes

Human-derived cultured neurons have the inherent drawback of a short life span. Human BD symptoms appear over months and years. This limits our electrophysiological measurements that can be acquired over weeks and cannot reflect the symptoms of the patients over years due to the different time scales. To solve this issue, we have developed a computational model that can be run over a large number of iterations. The cultured neurons change their properties over time gradually. Ion channels need to be expressed and modulated, and therefore, modifications in the currents are continuous and dynamic, and change slowly over time^52,53^. This should be reflected in a long-term computational model of the neurons. However, the simulated time-varying neuron should have the same statistical properties as the live measured neurons, projecting weeks of differentiation into thousands of simulation iterations, modeling years of neuronal functional, physiological properties.

To model this and to maintain continuous time-varying ion currents with the same statistical properties as the live neurons, we used a random walk algorithm. In this random walk, each ion channel is described as a stochastic process that follows a path that consists of a succession of random steps. The parameters of these steps were carefully chosen (see *Methods*) so that, in the end, after 10,000 iterations, this ion channel conductance will have the same statistical properties (using the KS test, see *Methods*) as the relevant measured ion channel. The cumulative distribution function of the recorded Na+, slow K+, and fast K+ currents for the three groups of neurons: control, LR, and NR, is plotted (Fig 4a-c; Supplementary Fig 4).

**Figure 4:**
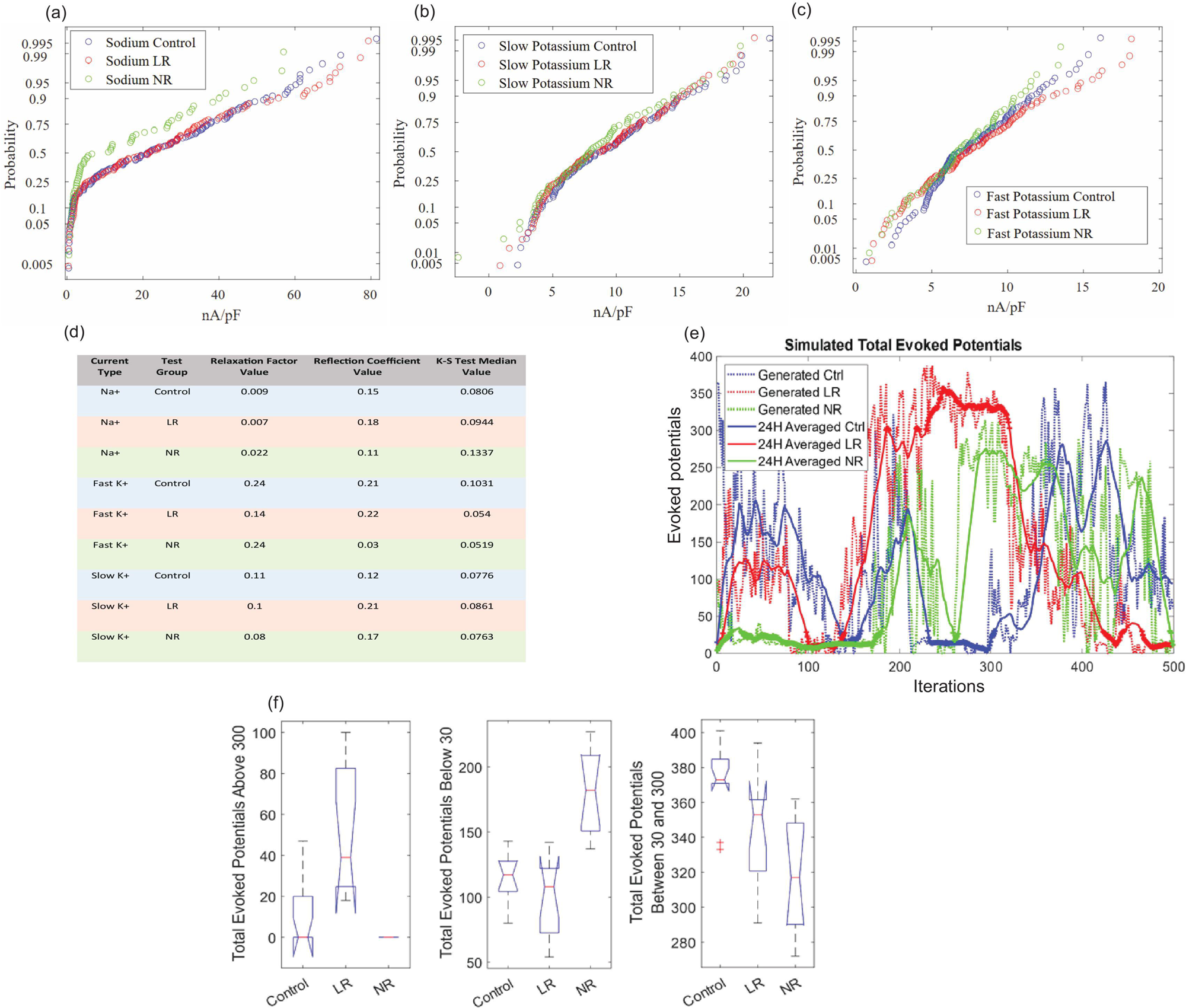
Long-term *in-silico* simulation of single-cell excitatory DG granule neurons incompletely captures the network dynamics associated with BD subtypes. The cumulative probability distributions of the amplitude of the (a) Na+, (b) slow K+, and (c) fast K+ normalized currents in control, LR, and NR DG excitatory neurons are plotted from the electrophysiological recordings^14,23,24^ from the same iPSC lines and cohort from which GABAergic neurons have been derived in this study (also see, *Methods*; Supplementary Fig 4). (d) The optimized configuration for the single cell model for each current type (Na+, fast K+, and slow K+) for each test group (control, LR, NR) (see *Methods*; Supplementary Fig 5). (e) The simulated number of evoked potentials over 500 iterations in control (blue), LR (red), and NR (blue) simulated neurons shows that the LR neuron exhibits a high activty state during these 500 iterations and the NR neuron exhibits a low activity state during these iterations. (f) The average excitability analysis results over 11 repeated simulations (500 iterations each) for the single-cell DG model show that the LR neuron produces over 300 APs in many iterations, mimicking a high activty-like state, while the NR neuron produces below 30 APs in many iterations, mimicking a low activity-like state.

Using the optimized parameters (Fig 4d; Supplementary Fig 5), the random walk model simulated long-term electrophysiological currents, generating 10,000 conductance samples per ion channel (Na+, slow K+, and fast K+) for a simulation of a control, LR, and NR neurons using our previously published model^24^ and with statistical parameters that are similar to the measured currents in our electrophysiological recordings. A 500-sample window was selected from the 10,000 samples with statistical parameters that are to the entire set of 10,000 samples. The single-cell model ran each iteration with a new ionic channel conductance according to the time-varying simulated currents. The excitability of the three types of simulated neurons (control, LR, and NR) was then assessed (Fig 4e). Analysis of the excitability showed that NR neurons tended to exhibit long periods of reduced excitability, reflecting possible low activity states of the neurons, while LR neurons tended to exhibit long periods of increased excitability, reflecting possible high activity states of the neurons. To quantify this, we defined two thresholds for hypo-excitability (<30 evoked potentials) and for hyperexcitability (>300 evoked potentials) (see *Methods*), and ran the model 11 times (randomly generating the 10,000 conductances each time) (Fig 4f). This quantification showed that LR neurons exhibit longer periods of hyperexcitability (high activity state) and NR neurons exhibit longer periods of hypoexcitability (low activity state). However, neither of the groups fully recapitulated both global hyperexcitability and global hypoexcitability states. The model in its current form did not explain the presence of both high activity and low activity clinical states in LR and NR patients.

### Integrating GABAergic neurons into a dentate gyrus network model enables both LR and NR networks to recapitulate essential features of mania and depression

We next created a biologically realistic multicellular network that resembles the rodent DG architecture, guided by experimentally reported cell-type ratios^9,54–57^ (Fig 5a). We used the single-cell model optimizations, and these were integrated into LR, NR, and control neuronal networks, where the ionic current shifts slowly over time, but their statistics remain similar to the measured ionic currents. The network was comprised of excitatory granule cells (GCs), excitatory hilar mossy cells (MCs) as the primary projection neurons, with an emphasis on the addition of basket cells (BCs) as GABAergic inhibitory neurons^58^ with changes in the BD neurons that were derived from the electrophysiological recordings (see *Methods*). The network featured 500 GCs, 25 BCs, 15 MCs, 6 hilar perforant path-associated cells (HIPPs), and 100 excitatory perforant path (PP) input cells, with connectivity following DG-specific principles^57,59^ (Fig 5b, Supplementary Tables 2,3). GCs formed AMPA and NMDA synapses with nearby MCs and BCs, projecting to a limited set of local BCs and interneurons to form feedback inhibitory loops. MCs provided excitatory synapses to local and distant GCs and interneurons, while BCs extended GABA-A inhibitory synapses to hundreds of GCs and other interneurons, all modeled with bi-exponential kinetics^54^ (Fig 5c, Supplementary Fig 6, Supplementary Tables 5-8).

**Figure 5:**
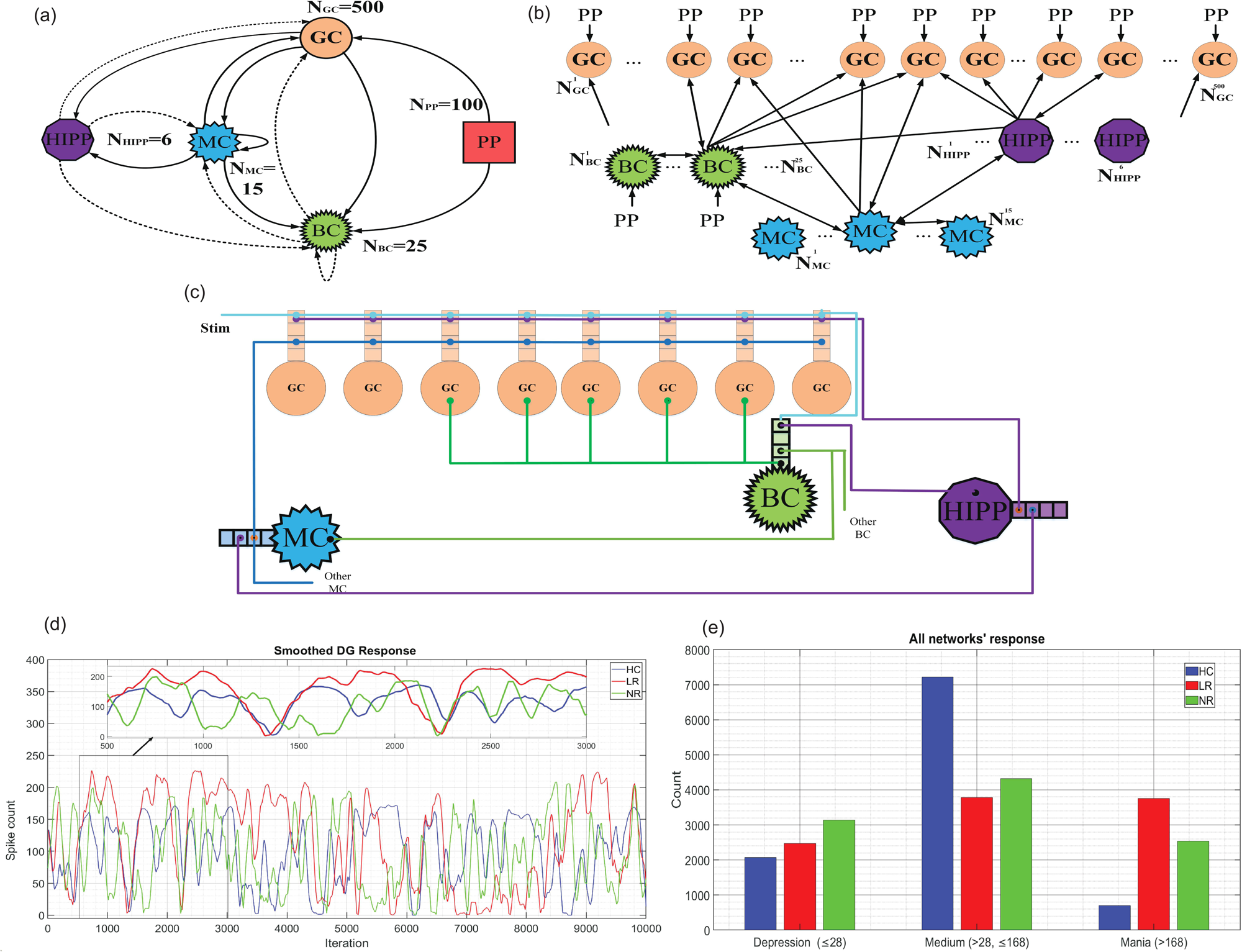
Integrating GABAergic neurons into a dentate gyrus network model enables both LR and NR networks to recapitulate essential features of mania and depression. A 10,000-iteration response of a full DG network of control (HC), LR, and NR. (a) Schematic representation of the biophysical (DG) network model. Granule cell (GC) models derived from LRs, NRs, and healthy controls (HCs) are integrated into a full DG circuit. The network consists of the perforant path (PP, red squares), basket cells (BC, green starbursts), hilar perforant path-associated cells (HIPP, purple octagons), and mossy cells (MC, blue bursts). Solid arrows represent excitatory synaptic connections, while dashed arrows indicate inhibitory projections. N is the population size for each cell type. This circuit diagram is adapted from our previous work ^42,43^. (b) Enhanced topological organization of the DG network, illustrating the distribution and connectivity among individual neurons and cell populations. The diagram highlights the layered structure and interplay between excitatory and inhibitory neuron types. (c) Detailed schematic of cellular connectivity within the DG model, depicting synaptic arrangements and pathways of information flow among GCs, BCs, HIPPs, and MCs. This circuit diagram is adapted from the previous study^59^. (d) The smoothed full DG circuit response simulations from all seven different networks with random synaptic connections for control (HC, blue), LR (red), and NR (green) (also see *Methods*; Supplementary Fig 7). The arrow shows a zoom in of the networks during 2500 iterations in the groups. (e) The average excitability and spike count analysis results over seven random network simulations for a DG circuit (10,000 iterations each) show that LR and NR full DG network spend more time in a low activity state compared to controls, and also more time in a high activty state compared to controls (also see, *Methods*; Supplementary Fig 7 and Supplementary Tables 5-8).

In our DG network simulations, we utilized 10,000 conductance parameters optimized from the single-cell model of excitatory neurons, representing 10,000 neuronal time units. We additionally changed BC different ion channel conductance parameters to be the mean measured currents of control, LR, and NR GABAergic neurons (according to our electrophysiological measurements shown in Fig 2). In this way, the conductance of the five channels that we measured to be different is summarized in Supplementary Table 5. The Na+ channel, rectifier K+, and A-type K+ conductance were scaled according to our electrophysiological measurements.

The DG network response was recorded across 500 GCs, and to ensure robustness, we analyzed responses across seven independent networks with randomized synaptic connections, preserving the original connection probabilities and synaptic weights (Supplementary Figs 7a & 7b). We found distinct response trajectories for control (HC), LR, and NR, when simulating 10,000 iterations representing 10,000 neuronal time units (see *Methods*) (Fig 5d provides the average smoothed spontaneous action potentials over the randomized 7 networks generated by the 3 types of networks: the control, LR, and NR networks). The networks average response (Supplementary Fig 7a & 7b shows each of seven network responses without and with smoothening, respectively, see *Methods*) revealed that both LR and NR networks spend more time in low activity states compared to controls (number of action potentials≤28, see *Methods*) and in high activity states compared to controls (number of action potentials >168, see *Methods*) (Fig 5e). Both LR and NR networks spend less time in the normal (Medium) range compared to the controls. These differences underscore how changes in neurophysiology shape a network that spends more time in low activity and high activity states. Importantly, only when these measured neurophysiological changes were applied to both excitatory and inhibitory neurons did the network start to cycle into these two states. Thus, our model captured the cyclical nature and dysregulation of E/I balance in BD network dynamics.

## DISCUSSION

Previous iPSC model-based investigations on excitatory neurons (hippocampal and cortical) revealed hyper-excitability in both BD subtypes (LR and NR), implying that altered neuronal firing is a primary feature of BD pathogenesis^9,14,60^. Interestingly, this hyperexcitability was neuronal-type dependent: While DG granule neurons derived from both BD LR and BD NR patients were hyperexcitable compared to controls, CA3 pyramidal neurons derived from BD LR patients were hyperexcitable, but BD NR CA3 pyramidal neurons were not ^9,14,22,23^. Also, when differentiating them as motor neurons, the three groups of control, LR, and NR did not exhibit significant electrophysiological differences. To better understand the role of inhibitory neurons, we established GABAergic-enriched neuronal cultures from iPSCs of healthy controls, LR, and NR BD patients. The electrophysiological measurements revealed hyper-excitability of LR GABAergic neurons, a possible compensation strategy against excitatory drive, but interestingly, NR GABAergic neurons were hypo-excitable compared to controls. In this work, we show that these alterations that we had observed in GABAergic activity of BD neurons are necessary for the neuronal network to develop states of global hyperexcitability (possibly mimicking a mania state) and global hypoexcitability (possible mimicking a depression state). It is important to keep in mind that high energy states also exist in depression^61,62^, yet studies have linked low activity with depression and high activity with mania^62–64^.

Transcriptomic profiling of iPSC-derived GABAergic neurons from BD patients reveals a complex molecular landscape that supports the divergent excitability patterns observed in LR and NR subtypes. Extensive DEGs were found between NR and control neurons which included downregulation of GABA receptor genes (*GABRA1, GABRR1, GABBR2*), GABAergic subtype (*VIP*), genes important for GABAergic neuron development (*DLX5, LHX2*), and for GABA transport and metabolism (*SLC6A11, ALDH5A1*), and Na+/K+ channel function genes (*KCNJ13, SLC5A12, KCNJ2, KCNE2*) compared to control neurons. These molecular abnormalities likely change the development of NR neurons and their neurophysiology as mature neurons, and can disrupt the inhibitory feedback required for network stability. In contrast, the transcriptome landscape of LR neurons relative to controls includes a more limited set of DEGs, with a notable dysregulation of metabolic pathway genes (*ENO4, GBE1, LDHA, MGAM2, PDK4, PPP1R14A, GLUD2, CH25H*) and upregulation of *GABRR1*. This suggests a metabolic reprogramming that might sustain the hyperexcitable state in LR neurons. Additionally, NR GABAergic neurons formed a distinct and separate cluster from control and LR neurons in a PCA plot (Supplementary Fig 3), aligning with their distinct molecular signature and altered neurophysiological properties.

To further test the premise of of dysregulated excitability in the network dynamics of BD subtypes, we first computationally simulated three types of neurons: LR, NR, and control. These were similar to our already published model^23,24^, however, we added more real-life scenarios by modeling changes in the ion channel conductance over time. To do this, we employed a random walk, a noise-driven system, to model how the electrical properties of BD neurons change over time. To keep the model realistic, the model possessed Gaussian white noise to mimic inherent stochastic variations in BD neurons while also setting statistical limits to avoid divergence of the model. We then implemented this in a single cell model comprising only excitatory neurons (LR, NR, and control). However, this model was limited and could only allow partial cycling of the network, with LR neurons predominantly showing hyperexcitable states and NR neurons exhibiting hypoexcitable states compared to the control neuron.

We then implemented a multicellular DG computational model^43,54^ integrating electrophysiological features of both excitatory and inhibitory neurons comprising 646 cells. The experimentally measured properties of the various Na+/K+ channels of patient-derived GABAergic neurons of LR, NR, and control were integrated into the model. The GCs comprised the major excitatory neurons, and BCs the GABAergic inhibitory neurons. . The model successfully recreated the hyper-excitable and hypo-excitable states seen in BD neurons and simulated long-term activity and energy state shifts (∼10,000 iterations). Importantly, it demonstrated that dysregulation of GABAergic excitability plays a major role in making BD networks unstable suggesting possible shift in the patients energy states. In contrast to previous conventional models that only present a static picture of brain activity, our dynamic stochastic model tracks the long-term progression of disease states. This is a significant step forward, allowing us to continuously monitor how neuron excitability changes over time and its effect on the overall network behavior. Additionally, to the best of our knowledge, our model is the first computational model that shows state shifts resembling mania and depression. Moreover, our model is more accurate and predictive because it uses empirical data from excitatory and inhibitory neurons of BD patients, all derived from the same iPSC lines.

In conclusion, we performed a detailed investigation of BD pathophysiology by integrating gene expression analysis, electrophysiological measurements, and computational modeling of patient-specific iPSC-derived neurons. This approach helped us shed light on BD progression and differential lithium response in BD subtypes. Furthermore, our study provides novel insights and addresses knowledge gaps regarding the role of inhibitory neuron dysfunction in BD and its effect on neural network stability. These insights might help to improve the development of targeted treatment methods for BD subtypes and explore other innovative therapeutic options.

## METHODS

### hiPSC lines of BD patients and control individuals

Nine iPSC lines comprising three controls (males) and six BD type I patients (three LRs and three NRs; all males) were used for this study as described and published previously^14^. Ethical approval was obtained at the Nova Scotia Health Authority, Canada. The iPSC experiments were performed following the relevant guidelines set by the institutional review board (IRB), University of Haifa, Israel, and approved by the IRB.

### Differentiation of hiPSCs into embryoid bodies (EBs)

We followed the recently published protocol for GABAergic neuron differentiation with minor modifications^44^. EBs were formed by mechanical dissociation of hiPSC colonies using dispase (07923, Stem Cell Technologies) and then plated onto low-adherence dishes in mTeSR medium with Rho kinase (ROCK) inhibitor. For EBs differentiation, floating EBs were treated with Dorsomorphin and A-83 (2 µM each) in Dulbecco’s modified Eagle’s medium (DMEM)/F12 (Invitrogen) plus knockout serum, Glutamax, beta-mercaptoethanol, and nonessential amino acid (NEAA) for gradually replacing mTESR media for 5 days. After 5 days, EB media was replaced with neural induction media (NIM) containing Smoothened agonist SAG (300nm), DMEM-F12 media containing N2 supplement, Non-essential amino acid (NEAA), and Glutamax.

### Generation of Neural Rosettes and Neural Progenitor Cells (NPCs)

To obtain Neural rosettes and NPCs, EBs were treated with above mentioned anti-caudalizing small molecule (SAG) and were plated onto Matrigel-coated plates in NIM for 2 weeks with alternate-day media changes. After 2 weeks, the neural rosettes were ready to be harvested when spheres showed neuronal extension and radial migration of cells. Neuronal Rosettes were manually picked and dissociated with Accutase (Chemicon) after 1 week and plated onto poly-L-ornithine and laminin-coated plates with GABAergic NPC media (DMEM/F12, N2 (1X), B27 (1X) (without retinoic acid), FGF2 (20 ng/ml), and laminin (1µg/ml)

### Differentiation of NPCs into GABAergic neurons

For GABAergic neuron differentiation, NPCs were plated on 6-well plates and 24-well plates with glass coverslips in GABAergic NPC media (as mentioned above). After 24-36 hours of seeding, the NPC media was replaced with Brainphys^TM^ media containing B27, Glutamax, Ascorbic acid (200 nM), cyclic AMP (cAMP; 500 mg/ml), laminin (1 µg/ml), BDNF (20 ng/ml), GDNF (20 ng/ml), and IGF-1 (20 ng/ml). The media was changed every other day, and experiments were performed at 21-28 days of differentiation.

### Lithium treatment of iPSC-derived GABAergic neurons

To chronically treat BD neurons with Li+, 1mM LiCl was added to the differentiation medium after 7 days in GABAergic neuronal differentiation media (composition mentioned above). The neurons were maintained for 4-5 weeks in culture, and the electrophysiology recordings were done as described below.

### Electrophysiology of iPSC-derived neurons

Whole-cell patch-clamp recordings were performed on cells after ∼3 weeks of differentiation. The bath contained artificial cerebrospinal fluid (ACSF) 10mM HEPES, 4mM KCl, 2 mM CaCl_2_, 1 mM MgCl_2_, 139 mM NaCl, 10 mM D-glucose (310 mOsm, pH 7.4). The recording micropipettes (tip resistance 8-10 Mohm) were filled with internal solution 130 mM K-gluconate, 6 mM KCl, 4 mM NaCl, 10 mM Na-HEPES, 0.2mM K-EGTA, 0.3 mM GTP, 2mM Mg-ATP, 0.2mM cAMP, 10mM D-glucose. The pH and osmolarity of the internal solution were brought close to physiological conditions (pH 7.3, 290–300 mOsmol). Amplification of signals was done with a Multiclamp 700B amplifier, and recording with Clampex 10.2 software (Axon Instruments). Signals were captured with Clampex at a sampling rate of 20 kHz and analyzed using Clampfit-10 and MATLAB (2018b, The MathWorks Inc., Natick, MA, 2000). All measurements were conducted at room temperature.

### Analysis of Electrophysiological Recordings

*(a) Sodium (Na+) and Potassium (K+) Currents:* To measure Na+ and K+ currents, neurons were held at -60 mV in voltage-clamp mode. We applied voltage steps ranging from −100 to +90 mV, each lasting 400 milliseconds. Na+ current was calculated by subtracting the Na+ current after stabilization from the lowest value of the inward Na+ current. Fast K+ current was measured by the maximum outward currents that appeared within a few milliseconds after a depolarization step. Slow K+ current was measured at the end of the 400-ms voltage step. All current values were normalized to each cell’s capacitance, which was measured using Clampex software. A one-way ANOVA was used to compare results between different groups.
*(b) Evoked action Potentials:* Neurons were held at -60 mV in current-clamp mode using a steady holding current. We then applied current injections in three pA steps (each lasting 400 ms), starting 12 pA below the holding current. A total of 38 increasing steps were applied^65^. The total evoked action potentials were counted across all 38 steps. Since the data were not normally distributed, we used the Wilcoxon signed-rank test (a non-parametric test) to compare groups
*(c) Spontaneous firing rate*: The spontaneous firing rate recordings were done by holding the neurons at −45mV in voltage-clamp mode. The frequency of the events for each cell was calculated by dividing the number of events by the duration of the recording (non-active cells were included with an event rate of 0 Hz). The mean rates and standard errors of the spontaneous firing rate of all the cells in each group were computed. Non-parametric statistical tests (Wilcoxon signed-rank) were performed for comparisons.

### Immunocytochemistry

Cells were fixed in 4% paraformaldehyde for 15 minutes and then blocked and permeabilized in PBS containing 0.1% Triton X-100 and 10% horse serum. Coverslips were incubated with the primary antibodies: rabbit anti-PAX6 (CST, mAb#60433, 1:250), mouse anti-NESTIN (CST, mAb#33475, 1:2000), mouse anti-SOX2 (SCBT, 365823,1:400) rabbit anti-NKX2.1/TTF-1 (Abcam, ab227652, 1:50), rabbit anti-VGLUT1 (Abcam, ab227805, 1:500) and Mouse anti-GABA (Abcam, ab86186, 1:400) in blocking solution overnight at 4°C, washed in PBS, incubated with DAPI (Abcam, ab228549, 1:2500) and the corresponding secondary antibodies for one hour at room temperature, washed, mounted on slides using Fluoromount G (Thermo Fisher), and dried overnight, protected from light. Fluorescence signals were detected using a confocal microscope, and images were processed with Zen and ImageJ.

### Bulk RNA Sequencing and analysis of GABAergic neurons

Total cellular RNA was extracted from approximately 3 million cells per sample in the third week of differentiation using the Zymo^TM^ RNA Clean and Concentrator Kit (Zymo Research, Cat no: R1013) according to the manufacturer’s instructions. The RNA samples’ quality and integrity were checked using the ND-1000 Nanodrop spectrophotometer (Thermofisher Scientific) and the Tape Station 2200 (Agilent). The RNA samples used for sequencing had RNA integrity number (RIN) values ≥ℒ8.5.

The library was prepared using the TruSeq Stranded Total RNA Prep Kit (Illumina) following the manufacturer’s instructions, and sequencing was performed on NovaseqX. Sequenced reads were quality-tested using FASTQC v0.11.5 and aligned to the hg38 human genome using the STAR aligner v2.78a. Mapping was carried out using default parameters, filtering non-canonical introns allowing up to 10 mismatches per read, and only keeping uniquely mapped reads. The expression levels of each gene were quantified by counting the number of reads that aligned to each exon or full-length transcript and normalized by its mean across all samples using HTseq v0.9.1. Differentially expressed genes (DEGs) were analysed for a false discovery rate (FDR) using the Benjamini-Hochberg^66^ rule for multiple hypothesis correction. Genes with an FDR (p-adjusted)L<L0.05 and |log2 fold changeL|L> 0.5 were included in the analysis (Supplementary Table 1), and the volcano plots were constructed for the same using custom MATLAB scripts. Gene Ontology (GO) enrichment and KEGG pathway analysis were performed using DAVID Bioinformatics Resources based on the *p*-value<0.05 (DEseq2), and network plots and pathways were constructed after analyzing for the FDR<0.05 for the enrichment of the pathways (Fig 3 and Supplementary Fig 2). A graphical representation of the gene network plots was constructed using nodes and edges using custom MATLAB scripts, where each node represented a pathway connected by the edges (lines). The size of each node is proportional to the number of DEGs. Principal component analysis (PCA) plots were plotted with DEGs *p*-value<0.05 (DEseq2) using custom MATLAB scripts (Supplementary Fig 2).

### Modeling single BD excitatory neurons with time-varying physiological alterations

To simulate 3 types of neurons: control, LR, and NR DG neurons, we used our previously published models^24^ with further modifications to make the computational neurons more similar to the live neurons. First, it was important to model changes over time that occur in live neurons. Hence, we assessed whether the data of measured currents follow a normal (Gaussian) distribution; we calculated skewness and kurtosis, normalized by their standard deviations, with values below 1.96 indicating normality. Our analysis revealed that fast K+ currents were normally distributed in the three groups: control, LR, and NR, while slow K+currents showed normality only in the LR and NR groups. Na+ channels tend to conduct low currents in all test groups and do not conduct currents with normal distribution in any of the groups (Supplementary Fig 4).

Further, we had to introduce differential changes in ion channel conductance, since in live neurons, large conductance changes cannot occur in a short time. However, these changes needed to occur such that the statistical properties of the computationally modeled ion channels are similar to the conductance changes that occur in the live neurons. For this, we incorporated changes in ion channel conductance as a random walk with an additional dissipation that does not allow the signal to diverge.

The following stochastic differential equation, based on the random walk, describes the channel conductance changes (sample i represents the ith time unit of the simulation run time).

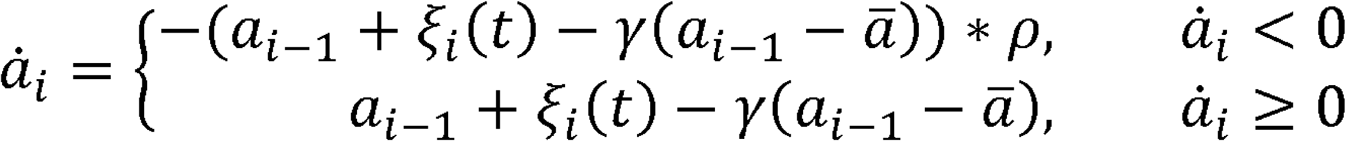

<(α – α̅)^2^ > = D/γ, variation of feature data

α̇_*i*_-represents "time unit" i of the simulated electrophysiological feature

α̅-represents the experimental mean of known feature data.

ξ_*i*_(*t*)-represents a random noise as a function of time for the predicted feature in a time unit, "i". We have used a white Gaussian noise with zero means and the variance that was proportional to the electrophysiological feature’s variance.

*γ* - represents the relaxation factor to stabilize the model.

*ρ* - represents the reflection coefficient to reflect negative data since the current values are not negative in a physiological scenario.

*D* - represents the variance of the noise (its spectral amplitude)

Membrane conductance of neurons varies due to various reasons, including stochastic closing and opening of ion channels even in the absence of synaptic inputs^52^, or due to fluctuations in synaptic inputs^67^. Additionally, variability can also arise due to molecular events such as changes in protein expression^53^, circadian rhythms^68^, or pathological processes^69^. This variability is supported by experimental studies of channel noise^52,70^ and stochastic adaptation models^71^. Prior publications have also shown that BD is a channelopathy^72^, and the iPSC-derived neurons are characterized by physiological instability^24^. Hence, to capture the temporal dynamics of neuronal conductance in BD, we modeled it as a time-varying parameter.

The reflection coefficients configure the amount of mirroring the model should use on negative currents. For the statistics of the simulated ionic currents to be the same as the statistics of the recorded electrophysiological values, we employed nested loops iterating through parameter value ranges, empirically selecting those that minimize the statistical differences between the measured and the statistics of the simulated currents. This was assessed via the Two-sample Kolmogorov-Smirnov (KS) test over 100 simulations for each ion channel conductance. The KS test, testing the null hypothesis of identical continuous distributions at 5% significance, returned "1" for rejection and "0" otherwise. The algorithm identified parameter values yielding the fewest rejections, indicating the closest match to physiological data, with median KS statistics recorded for future reference. This process generated a three-dimensional grid (parameters vs. rejection counts), revealing a minimum point for each ion channel ^24^conductance. Nine parameter sets (three ionic currents × three groups) were optimized, and these are detailed in Supplementary Figure 5 and Figure 4d.

#### Electrophysiology parameters fitting in the single-cell computational model

The model used measured electrophysiology ionic channel currents to determine differences in features like action potentials and excitability between LR and NR BD patients. Python scripts using SciPy.signal and NumPy packages analyzed action potentials and excitability features. The computational model performed simulations of neuronal network health and disease states in the NEURON simulation environment using parameters such as sodium and potassium ion channel capacitance and conductance (Na, Ka, Km, Kdr, Kcnc).

### Dentate Gyrus Network

The DG network model was developed to investigate hippocampal function and BD-associated dynamics, replicating rodent DG microcircuitry^42,54,73^. GCs, the principal projection neurons, received excitatory input via perforant paths (PP) synapses, with axons projecting to CA3, while interneurons mediated indirect feedback loops (Fig 5a and b). The network, constructed in NEURON 8.x (described below) using cell-type ratios included 500 GCs, 25 BCs, 15 MCs, 6 HIPPs, and 100 perforant paths (PP) inputs, with AMPA, NMDA, and GABA-A synapses modeled using bi-exponential kinetics^57,59^ (Fig 5c, Supplementary Fig 6, and Supplementary Tables 2, 5, and 8).

### Multicellular network connectivity and model construction for BD and control

Single-cell models were embedded into a multicompartmental network, with somas and dendrites segmented based on anatomical data^59^ (Fig 5b, Supplementary Table 6). GCs formed excitatory synapses with local BCs and MCs, while MCs projected to GCs and interneurons, and BCs provided broad inhibitory input. Pathological adaptations simulated BD states (control, LR, NR) by adjusting conductance parameters, validated against hyperexcitability and epileptogenesis studies^54,55^. The model integrated 10,000 conductance parameters optimized from the single-cell simulations described in the previous section (Supplementary Fig 5 and Fig 4d), fine-tuning BC conductances and scaling sodium, rectifier potassium, and A-type potassium channels via grid search (Supplementary Tables 5-8) with the same statistics as the live electrophysiological neuronal recordings. Seven independent networks with randomized synaptic connections that preserved original probabilities and weights were simulated (Supplementary Fig 7a and 7b). The 10,000 iterations simulated capturing long-term dynamics (see Supplementary Tables 5-8, Fig 5c and d, Supplementary Fig 7) of control vs. BD DG networks.

### Simulation and analysis of the model framework

All simulations were executed using the NEURON 8.x platform^41,74^. Single-cell biophysical properties and synaptic mechanisms were programmed in NMODL, while network assembly, stimulation protocols, and data acquisition were managed through NEURON’s HOC scripting language. The NEURON GUI facilitated interactive development to confirm cell responses and connectivity, though final runs were performed headlessly on a computing cluster. Simulations were carried out at 35°C with a fixed time-step integration (dt=0.025 ms) to precisely track action potential dynamics. Network simulations, spanning 5–10 seconds of biological time, evaluated steady state and transient behaviors, avoiding NEURON’s Code variable-step integrator to ensure uniform numerical stability across trials. Each run included a 2-second equilibration period to stabilize network activity, followed by 3 seconds of data recording. Baseline activity was maintained by delivering independent Poisson spike trains (10 Hz) to each granule cell via NetStim/NetCon, mimicking cortical input. Controlled experimental stimuli were introduced using NEURON’s IClamp and VecStim objects to apply somatic current injections or synaptic barrages to selected cells. Random seeds were set for reproducibility, and the code (including mod files and HOC scripts) was designed for seamless switching between LR, NR, control parameter sets, and physiological or pathological configurations.

To assess network behavior under different experimental conditions, somatic membrane potentials were recorded from representative GCs within the DG model. Key electrophysiological metrics, primarily the number of evoked spikes, were extracted from voltage traces using custom Python scripts. Identical stimulation protocols were applied across all groups-LRs, NRs, and controls to ensure consistent comparisons.

## ACKNOWLEDGMENTS

Prof. Shani Stern acknowledges the Zuckerman STEM leadership program and Israel Science Foundation grants - 1994/21 and 3252 /21 for supporting the research. Further, we acknowledge the fellowships provided by the Graduate Studies Authority, Bloom Graduate School, University of Haifa, Israel, for supporting our research. The schematic images in the manuscript were created with Biorender.com.

## DATA AND CODE AVAILABILITY

The data that support the findings of this study are available upon request from the authors. The codes for electrophysiology data analysis and neuron network modeling are available upon reasonable request from the authors.

## Supplementary Materials legends

**Supplementary Fig 1: Lithium treatment does not change the excitability of LR and NR GABAergic neurons.** The example traces for evoked action potentials of LR GABAergic neurons are shown in (a) LR with Li+ treatment and (b) LR without treatment. (d) The average evoked potentials with no significant change in the number of APs for LR with and without Li+ treatment. Similarly, the example traces for evoked action potentials are shown in e) NR neurons with Li+ treatment and (f) LR (red) and NR neurons without Li+ treatment (green). (g) The average evoked potentials with no significant change in the number of APs for NR with and without Li+ treatment. *Data are presented as mean ±SEM*. \**p*<0.05, \*\**p*<0.01, \*\*\**p*<0.001.

**Supplementary Fig 2: KEGG enrichment pathways after transcriptomic analysis of control, LR, and NR neurons.** The KEGG pathway enrichment analysis for the differentially expressed genes *(p*-value <0.05) between (a) LR vs control, (b) NR vs control, (c) NR vs LR, and (d) BD vs control conditions. The size of each pathway is proportional to the number of DEGs. FDR cut-off at <0.05.

**Supplementary Fig 3:** PCA plot of the control, LR, and NR groups (with replicates) using the differentially expressed genes (*p*<0.05, DEseq2) shows a further segregated clustering of NR groups from control and LR groups.

**Supplementary Fig 4: Distribution properties of Na+/K+ conductance from control, LR, and NR groups.** The distribution of measured electrophysiological currents for (a) Na+, (b) fast K+, and (c) slow K+ channels in control, LR, and NR excitatory neurons are plotted from the electrophysiological recordings of excitatory hippocampal neurons from the previous study^14,23,24^ from the same iPSC lines and cohort from which GABAergic neurons have been derived in this study (also see, *Methods*; Supplementary Fig 4)

**Supplementary Fig 5:** Optimization of Na+/K+ conductance of excitatory hippocampal neurons via random walk for the construction of a single cell model. a) Sodium (Na+) optimization process using Two-sample Kolmogorov-Smirnov (KS) test statistics. (b) The number of rejected null hypotheses for the Na+ optimization process. (c) Fast potassium (K+) optimization process using KS test statistics. (d) The number of rejected null hypotheses for the Fast K+ optimization process. (e) Slow K+ optimization process using KS test statistics. (f) The number of rejected null hypotheses for the slow K+ optimization process (also see *Methods* Fig 4d, Supplementary Fig 4)

**Supplementary Fig 6: Synaptic weights between source and target cell types in the dentate gyrus (DG) network**. (a) Perforant path (PP) projections to all other cell types. (b) Granule cell (GC) projections to target cells. (c) Basket cell (BC) outputs to other cell types. (d) Mossy cell (MC) projections. (e) Hilar perforant path-associated cell (HIPP) outputs. (f) Combined synaptic weight matrix showing all source-to-target cell connections.

**Supplementary Fig 7:** Seven independent networks with randomized synaptic connections that preserved original probabilities and weights were simulated. (a) Non-smoothed network response. (b) Smoothed network response

**Supplementary Table 1**: List of total DEGs between LR vs control, NR vs control, NR vs LR, and BD vs control conditions with their *p*-values, FDR (p-adjusted values), and Log2 fold changes.

**Supplementary Table 2:** Synaptic Weight Matrix Between Cell Types in the Dentate Gyrus (DG) Network

**Supplementary Table 3:** Neuron Section Dimensions and Segmentation Details Across Cell Types

**Supplementary Table 4**: Granule Cell Conductance Parameters within HC, LR, and NR **Supplementary Table 5**: Basket Cell Conductance Parameters within HC, LR, and NR **Supplementary Table 6**: Mossy Cell Conductance Parameters within HC, LR, and NR **Supplementary Table 7**: Mossy Cell Conductance Parameters within HC, LR, and NR

Supplementary **Table 8** : Synaptic Parameters for Granule Cell (GC), Basket Cell (BC), Mossy Cell (MC), and HIPP Interneurons.

