## Supplementary Figures1-7 for "Divergent Excitability of GABAergic Neurons Derived from Bipolar Disorder Patients Shapes Energy Shifts of Network Dynamics, possibly mimicking mania and depression"

### Supplementary Fig 1

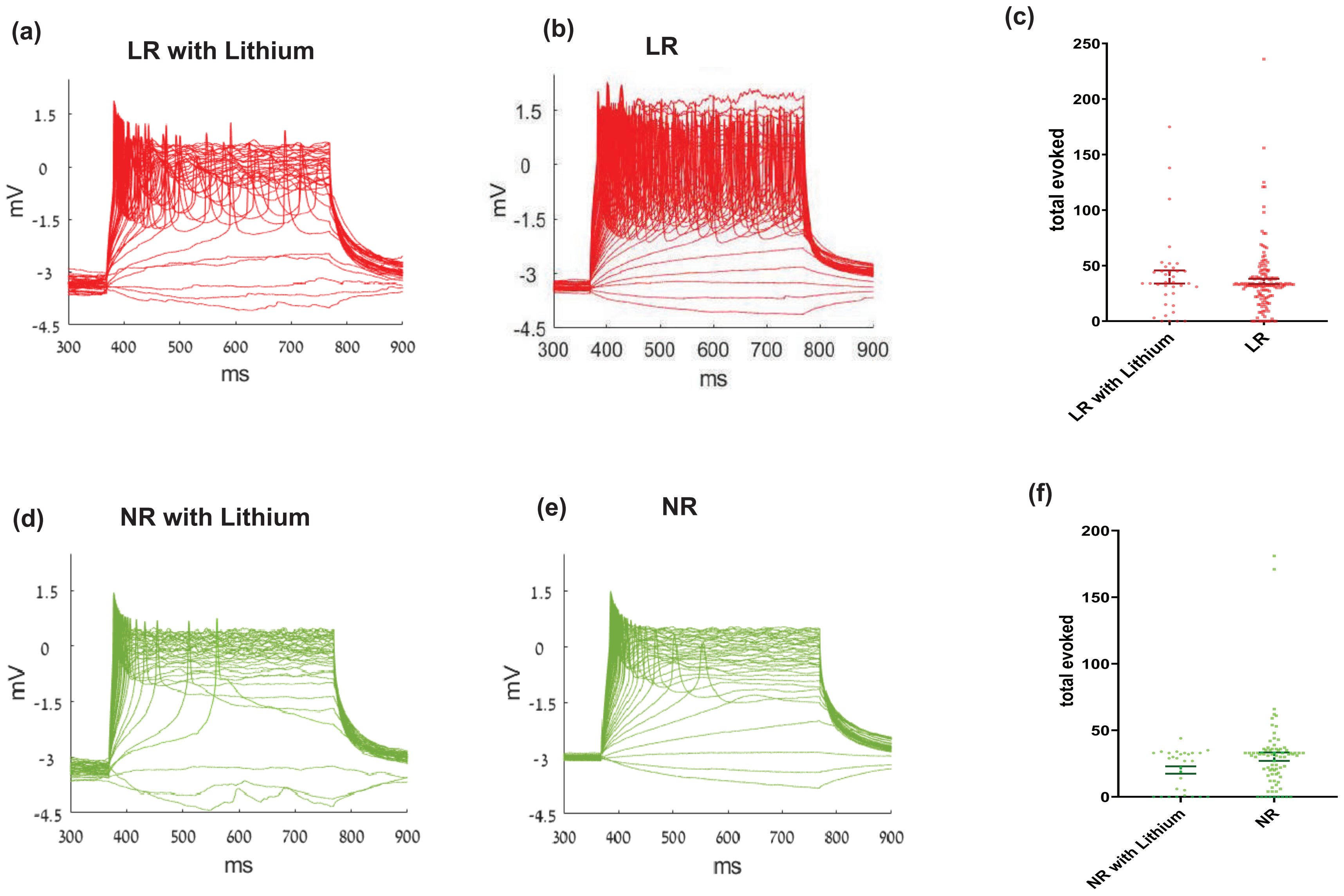

Supplementary Fig 2

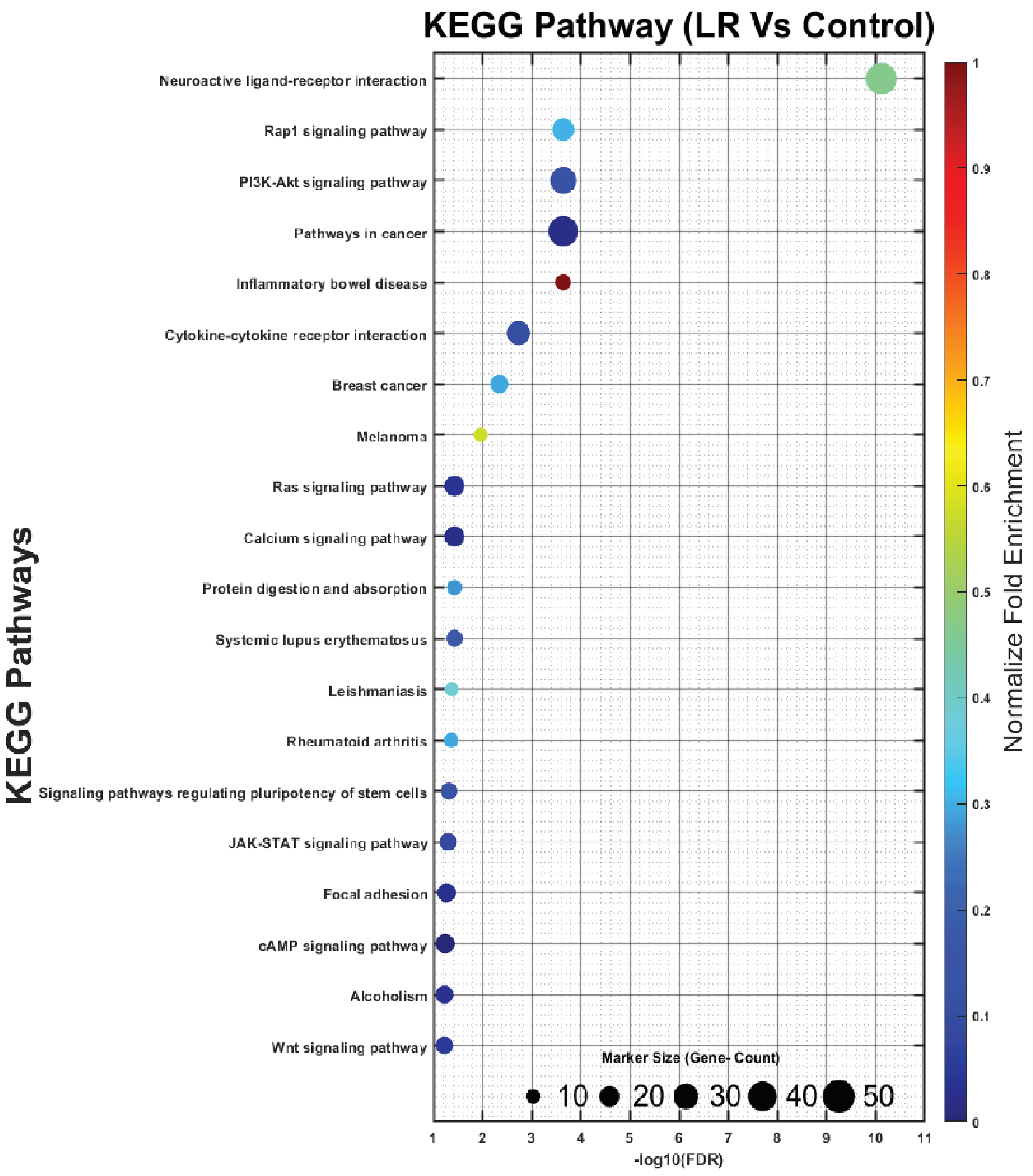


(a)

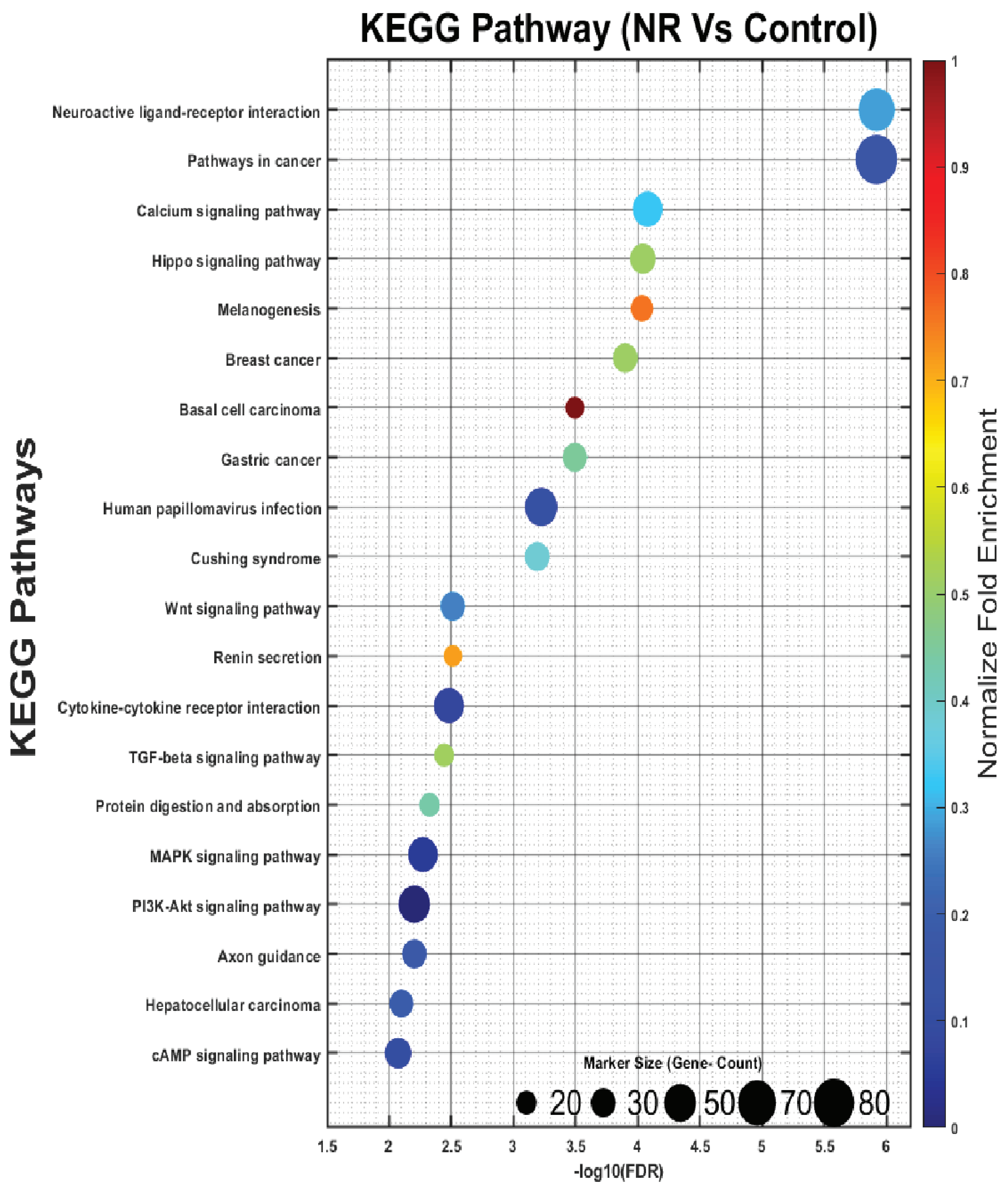


(b)

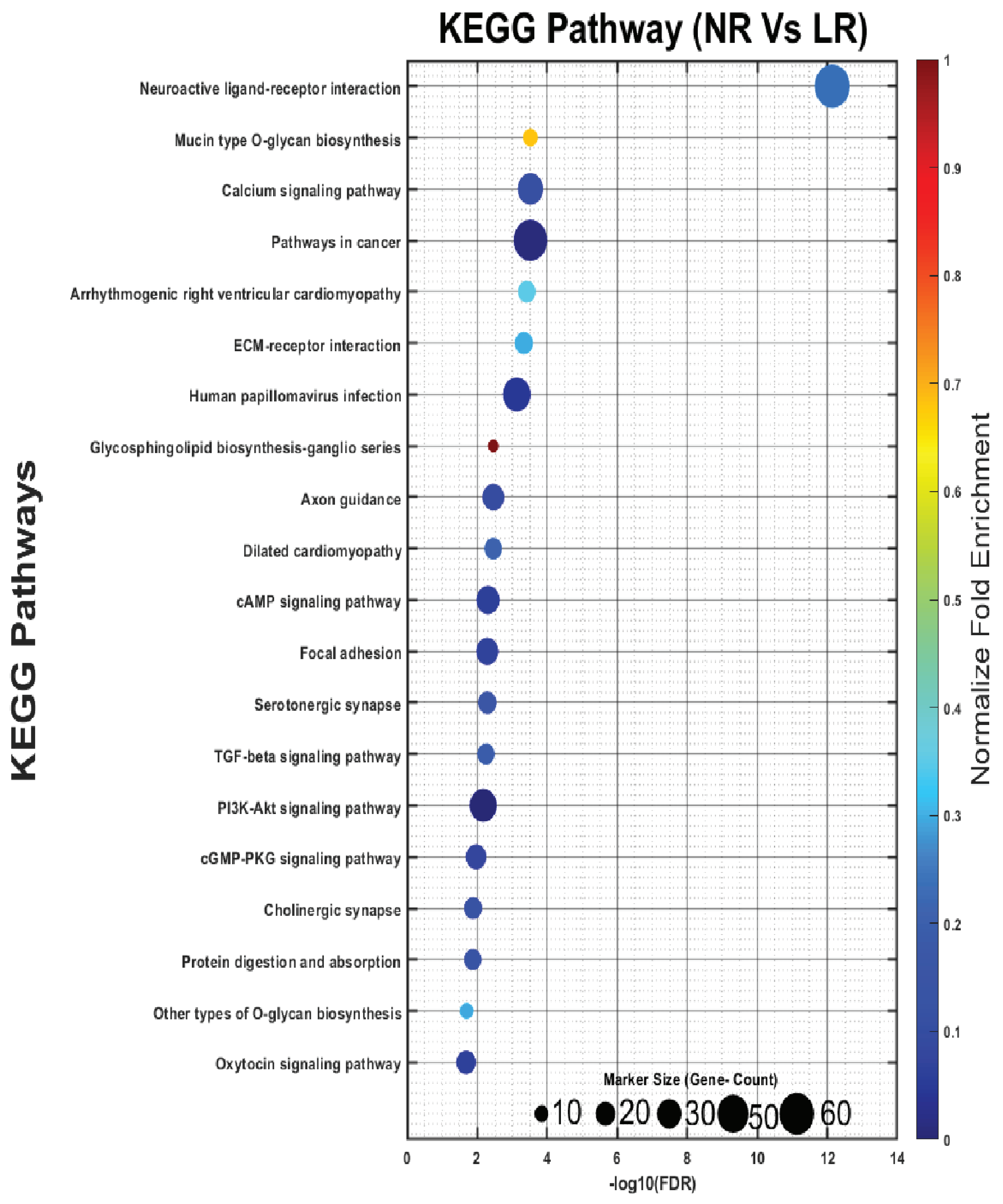


(c)

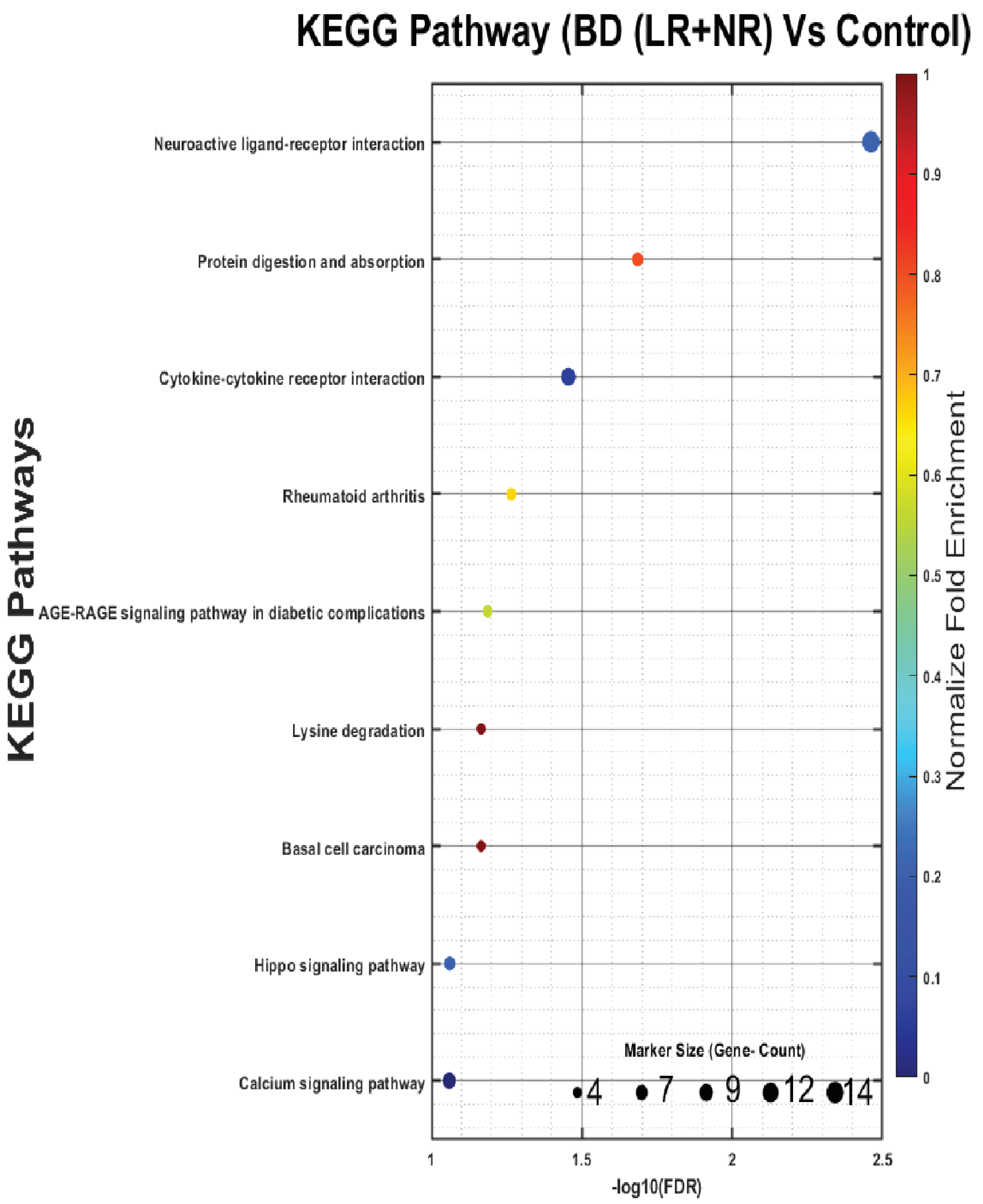


(d)

**Supplementary Fig 3**

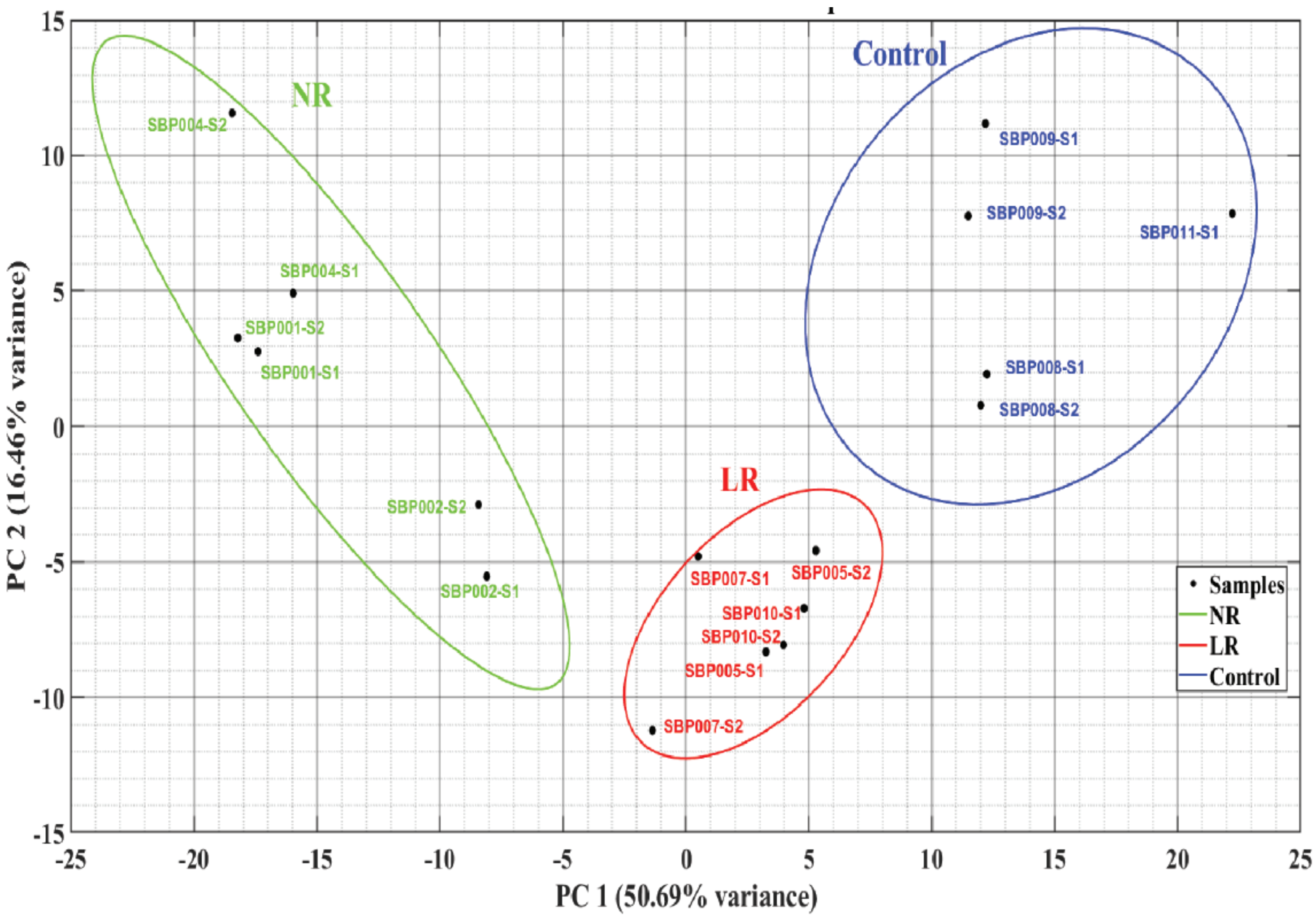

Supplementary Fig 4

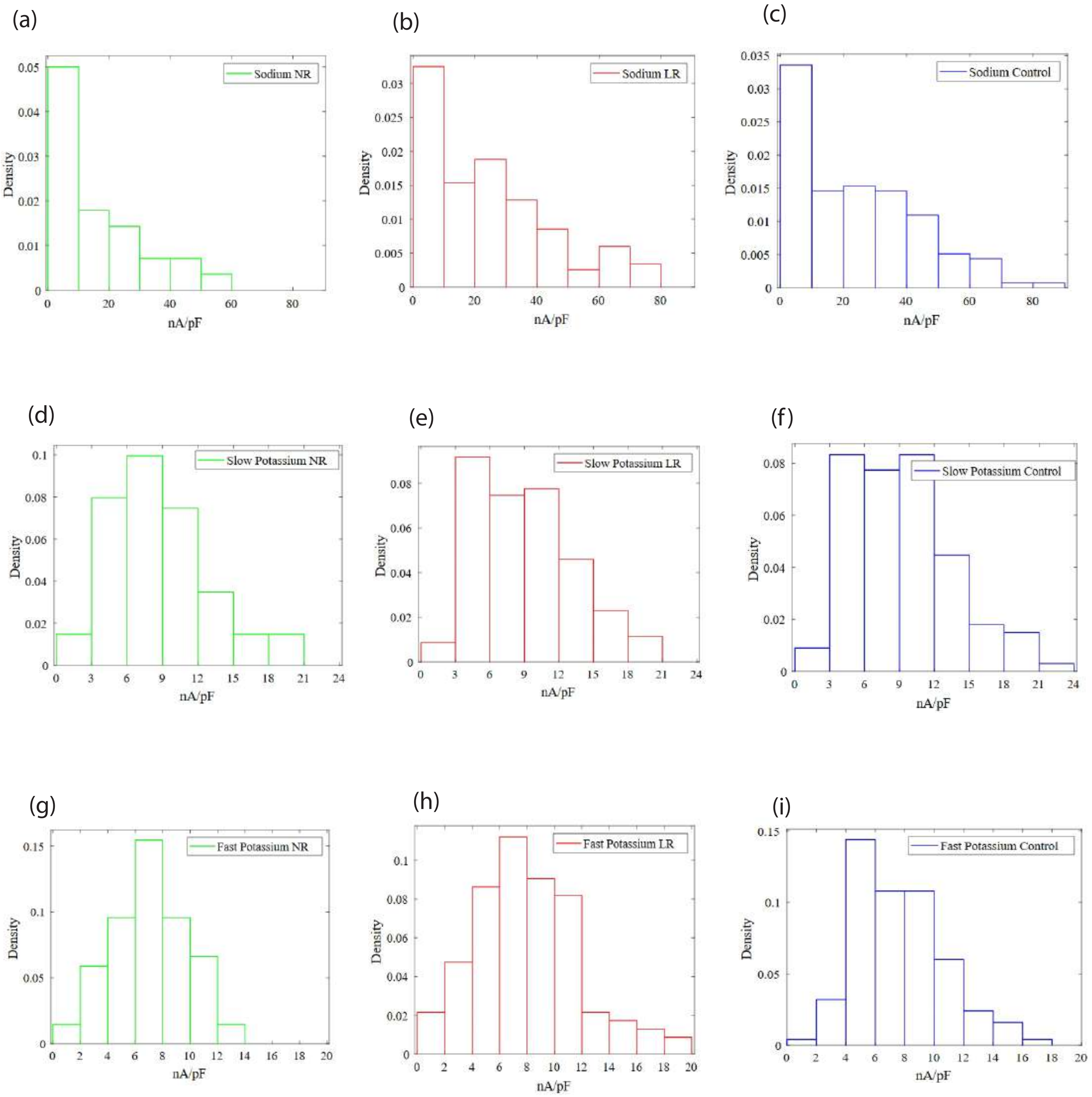

Supplementary Fig 5

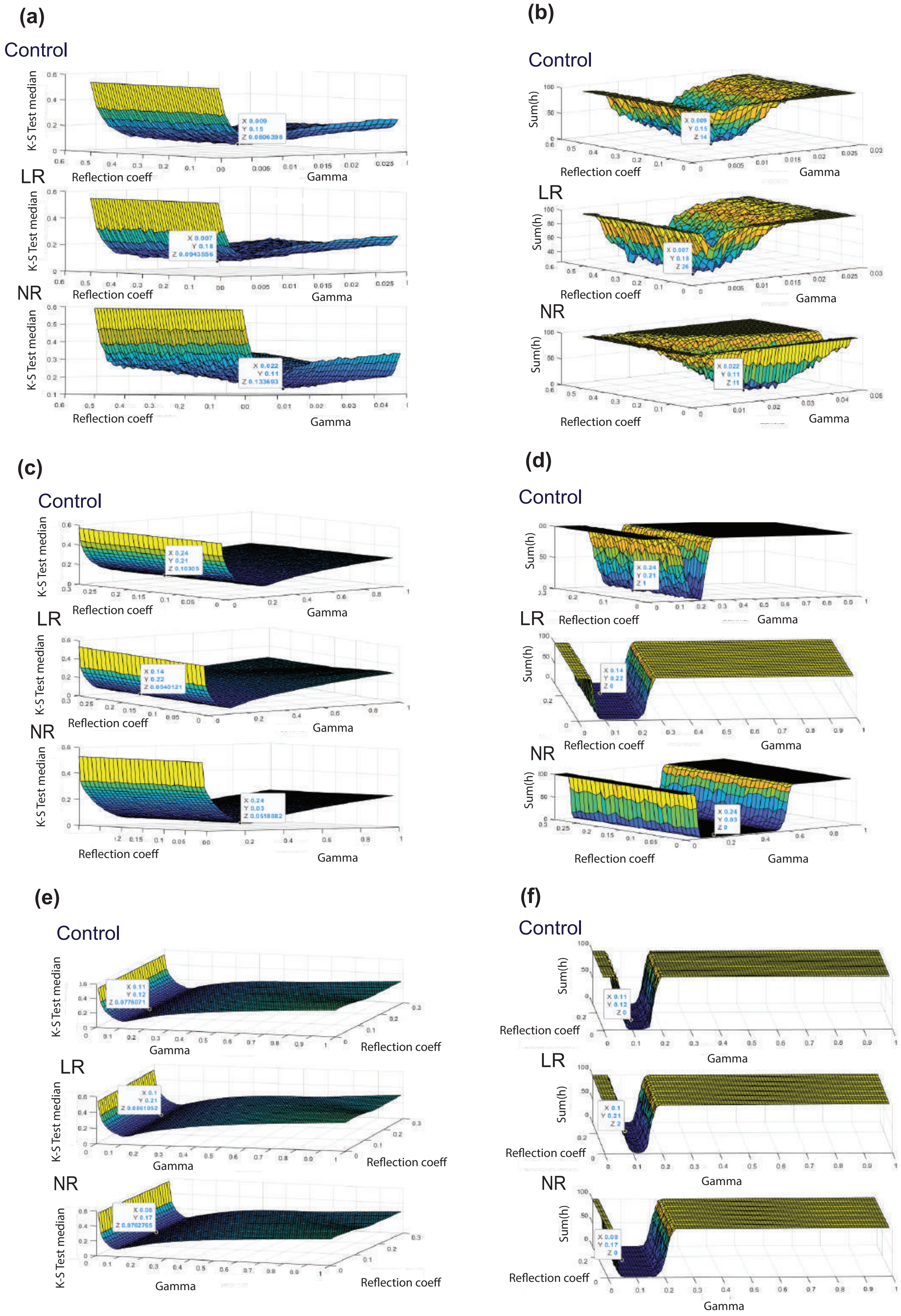

**Supplementary Fig 6**

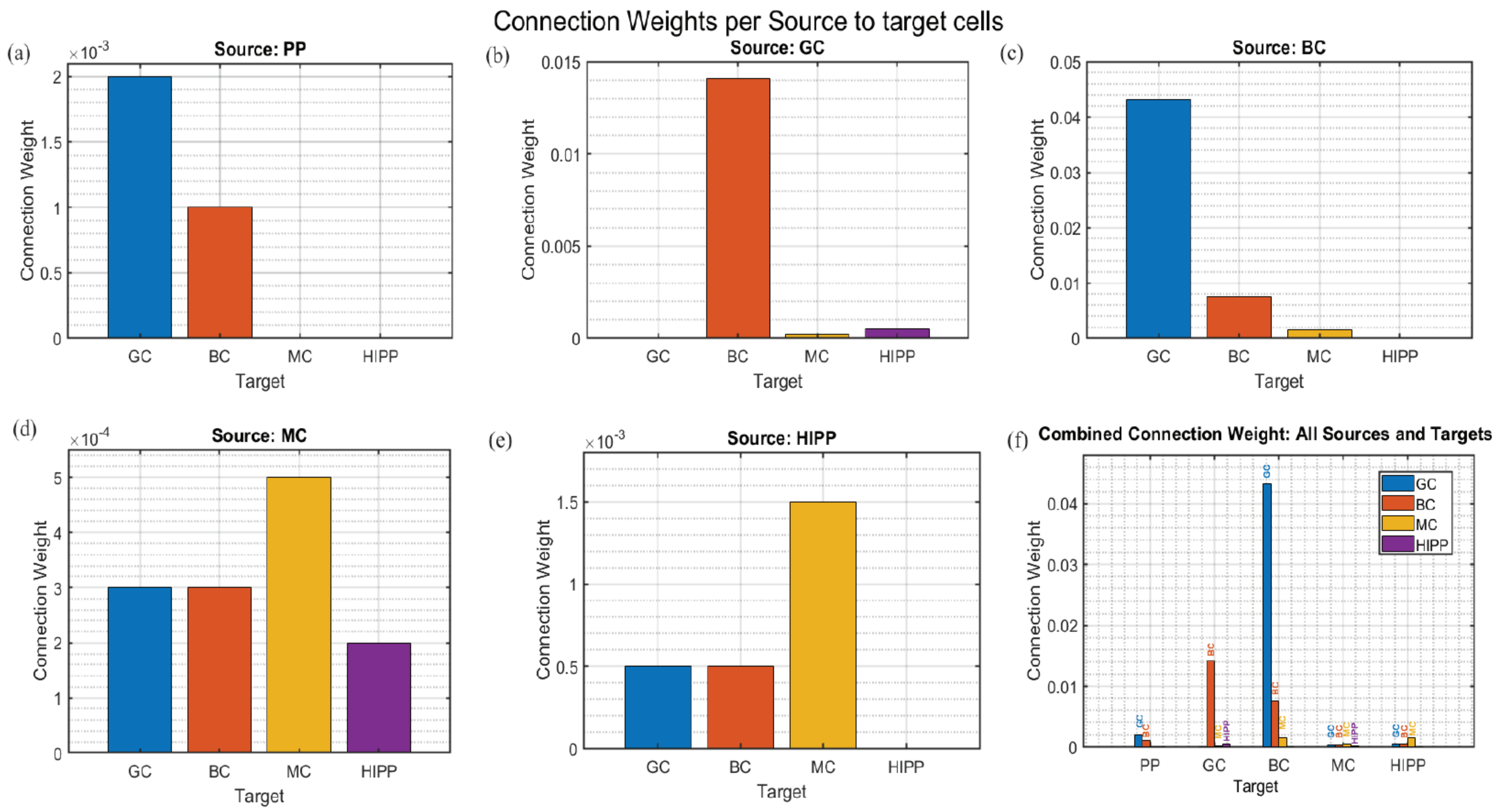

**Supplementary Fig 7**

**(a)**

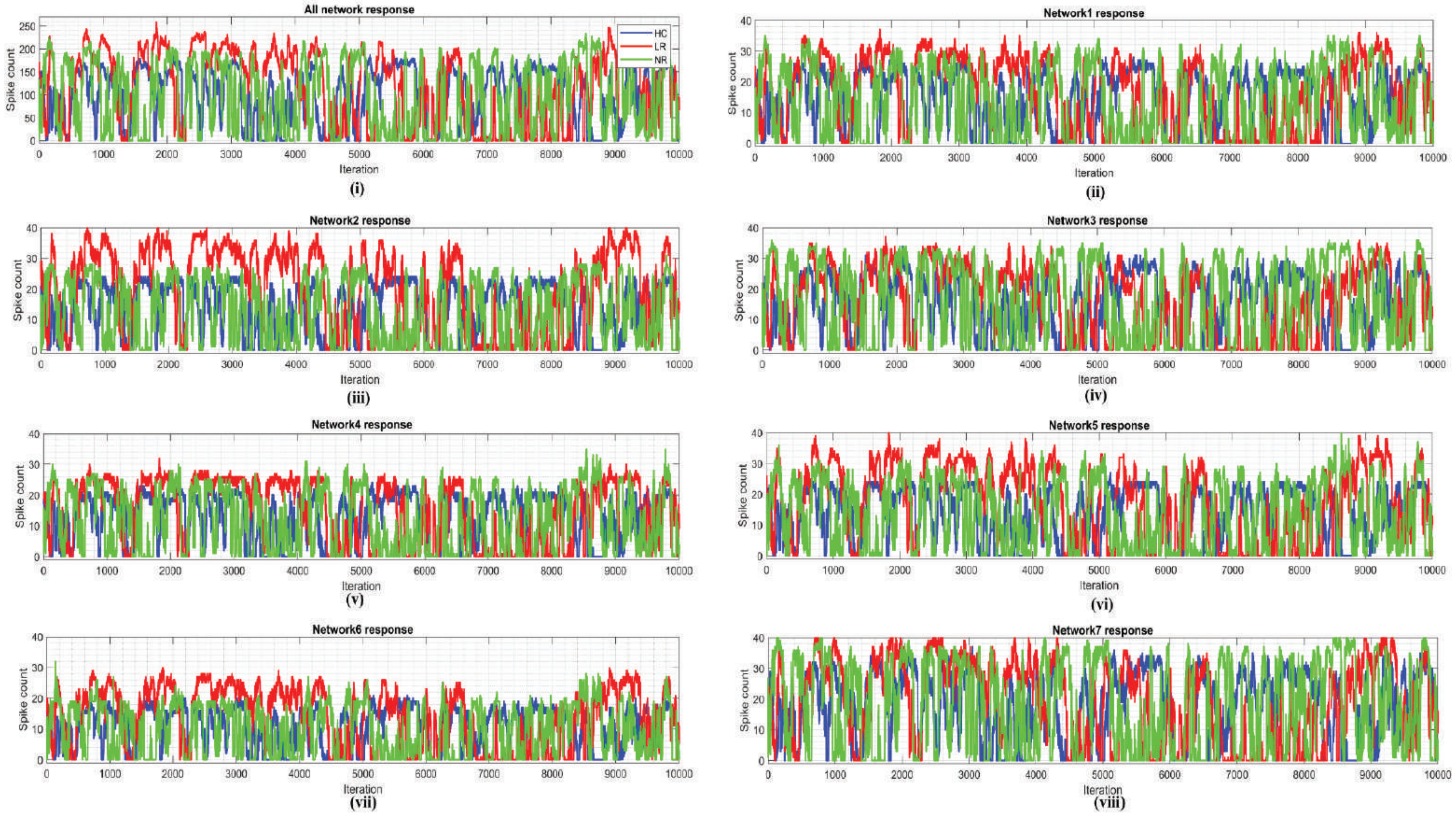


**(b)**

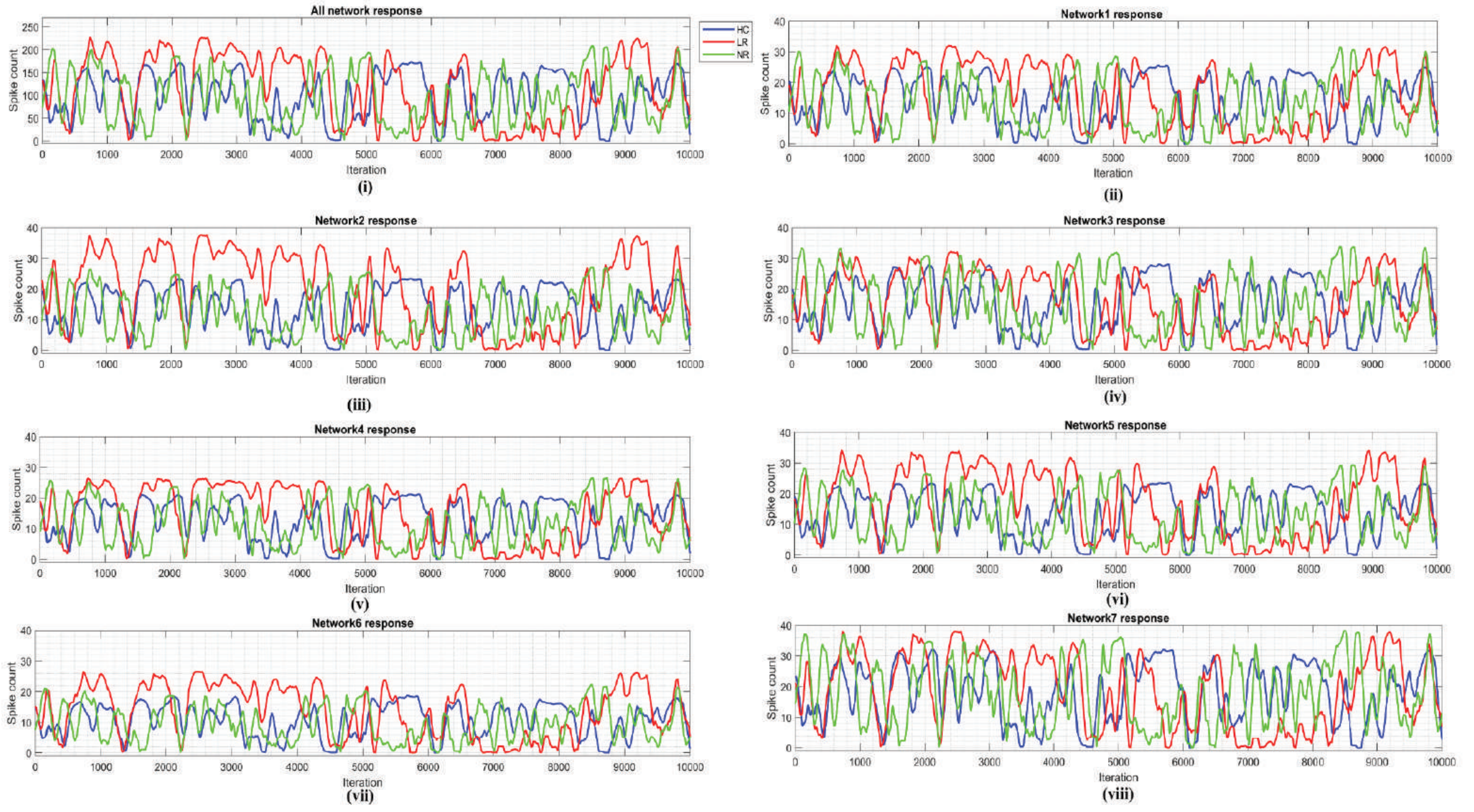
