## Supplementary Tables 2-8 for "Divergent Excitability of GABAergic Neurons Derived from Bipolar Disorder Patients Shapes Energy Shifts of Network Dynamics, possibly mimicking mania and depression"

Supplementary Table 2: Synaptic Weight Matrix Between Cell Types in the Dentate Gyrus (DG) Network

| **From \ To** | **GC** | **BC** | **MC** | **HIPP** |
| --- | --- | --- | --- | --- |
| **PP** | 0.0020 | 0.0010 | 0 | 0 |
| **GC** | 0 | 0.0141 | 0.0002 | 0.0005 |
| **BC** | 0.0048 | 0.0076 | 0.0015 | 0 |
| **MC** | 0.0003 | 0.0003 | 0.0005 | 0.0002 |
| **HIPP** | 0.0005 | 0.0005 | 0.0015 | 0 |

Supplementary Table 3: Neuron Section Dimensions and Segmentation Details Across Cell Types

| **Cell Type** | **Section** | **Index** | **Length (µm)** | **Diameter (µm)** | **Description** |
| --- | --- | --- | --- | --- | --- |
| **Granule Cell** | soma | — | 16.8 × 0.885 = 14.86 | 16.8 × 0.885 = 14.86 | Soma size |
|  | gcdend1 | 0 | 50 × 0.885 = 44.25 | 3 × 0.885 = 2.66 | Dendrite branch 1 |
|  | gcdend1 | 1 | 150 × 0.885 = 132.75 | 3 × 0.885 = 2.66 |  |
|  | gcdend1 | 2 | 150 × 0.885 = 132.75 | 3 × 0.885 = 2.66 |  |
|  | gcdend1 | 3 | 150 × 0.885 = 132.75 | 3 × 0.885 = 2.66 |  |
|  | gcdend2 | 0 | 50 × 0.885 = 44.25 | 3 × 0.885 = 2.66 | Dendrite branch 2 |
|  | gcdend2 | 1 | 150 × 0.885 = 132.75 | 3 × 0.885 = 2.66 |  |
|  | gcdend2 | 2 | 150 × 0.885 = 132.75 | 3 × 0.885 = 2.66 |  |
|  | gcdend2 | 3 | 150 × 0.885 = 132.75 | 3 × 0.885 = 2.66 |  |
| **Basket Cell** | soma | — | 20 | 20 | Soma size |
|  | bcdend1 | 0 | 75 | 4 | Apical dendrite |
|  | bcdend1 | 1 | 75 | 3 | Apical dendrite |
|  | bcdend1 | 2 | 75 | 2 | Apical dendrite |
|  | bcdend1 | 3 | 75 | 1 | Apical dendrite |
|  | bcdend2 | 0 | 75 | 4 | Apical dendrite |
|  | bcdend2 | 1 | 75 | 3 | Apical dendrite |
|  | bcdend2 | 2 | 75 | 2 | Apical dendrite |
|  | bcdend2 | 3 | 75 | 1 | Apical dendrite |
|  | bcdend3 | 0 | 50 | 4 | Basal dendrite |
|  | bcdend3 | 1 | 50 | 3 | Basal dendrite |
|  | bcdend3 | 2 | 50 | 2 | Basal dendrite |
|  | bcdend3 | 3 | 50 | 1 | Basal dendrite |
|  | bcdend4 | 0 | 50 | 4 | Basal dendrite |
|  | bcdend4 | 1 | 50 | 3 | Basal dendrite |
|  | bcdend4 | 2 | 50 | 2 | Basal dendrite |
|  | bcdend4 | 3 | 50 | 1 | Basal dendrite |
| **Mossy Cell** | soma | — | 20 | 10 | Soma size |
|  | mcdend1 | 0 | 50 | 5.78 | Dendrite branch 1 |
|  | mcdend1 | 1 | 50 | 4 |  |
|  | mcdend1 | 2 | 50 | 2.5 |  |
|  | mcdend1 | 3 | 50 | 1 |  |
|  | mcdend2 | 0 | 50 | 5.78 | Dendrite branch 2 |
|  | mcdend2 | 1 | 50 | 4 |  |
|  | mcdend2 | 2 | 50 | 2.5 |  |
|  | mcdend2 | 3 | 50 | 1 |  |
|  | mcdend3 | 0 | 50 | 5.78 | Dendrite branch 3 |
|  | mcdend3 | 1 | 50 | 4 |  |
|  | mcdend3 | 2 | 50 | 2.5 |  |
|  | mcdend3 | 3 | 50 | 1 |  |
|  | mcdend4 | 0 | 50 | 5.78 | Dendrite branch 4 |
|  | mcdend4 | 1 | 50 | 4 |  |
|  | mcdend4 | 2 | 50 | 2.5 |  |
|  | mcdend4 | 3 | 50 | 1 |  |
| **HIPP Interneuron** | soma | — | 20 | 10 | Soma size |
|  | hcdend1 | 0 | 75 | 3 | Dendrite branch 1 |
|  | hcdend1 | 1 | 75 | 2 |  |
|  | hcdend1 | 2 | 75 | 1 |  |
|  | hcdend2 | 0 | 75 | 3 | Dendrite branch 2 |
|  | hcdend2 | 1 | 75 | 2 |  |
|  | hcdend2 | 2 | 75 | 1 |  |
|  | hcdend3 | 0 | 50 | 3 | Dendrite branch 3 |
|  | hcdend3 | 1 | 50 | 2 |  |
|  | hcdend3 | 2 | 50 | 1 |  |
|  | hcdend4 | 0 | 50 | 3 | Dendrite branch 4 |
|  | hcdend4 | 1 | 50 | 2 |  |
|  | hcdend4 | 2 | 50 | 1 |  |

Supplementary Table 4: Granule Cell Conductance Parameters within HC, LR, and NR

| **Section** | **Mechanism** | **Parameter** | **Base Value / Expression** | **Scaling Factor (HC, LR, NR)** | **HC Value** | **LR Value** | **NR Value** |
| --- | --- | --- | --- | --- | --- | --- | --- |
| all | ichan2 | ggabaa_ichan2 | 7.22E-06 | HC=1, LR=1, NR=1 | 0.00000722 | 0.00000722 | 0.00000722 |
| **soma** | ichan2 | gnatbar_ichan2 | 0.12 | HC=1, LR=1.03, NR= 0.7878 | **0.12** | **0.1236** | **≈ 0.094545** |
| soma | ichan2 | gkfbar_ichan2 | 0.016 | HC=1, LR=1.15, NR=1.3 | **0.016** | **0.0184** | **0.0208** |
| soma | ichan2 | gksbar_ichan2 | 0.006 | HC=1, LR=1.15, NR=1.3 | **0.006** | **0.0069** | **0.0078** |
| soma | ichan2 | gl_ichan2 | 2.27E-05 | HC=1, LR=1, NR=1 | 2.27E-05 | 2.27E-05 | 2.27E-05 |
| soma | ka | gkabar_ka | 0.012 | HC=1, LR=1.2 , NR=1.3 | **0.012** | **0.0144** | **0.0156** |
| soma | km | gbar_km | 0.001 | HC=1, LR=1.15, NR=1.3 | **0.001** | **0.00115** | **0.0013** |
| soma | nca | gncabar_nca | 0.002 | HC=1, LR=1, NR=1 | 0.002 | 0.002 | 0.002 |
| soma | lca | glcabar_lca | 0.005 | HC=1, LR=1, NR=1 | 0.005 | 0.005 | 0.005 |
| soma | tca | gcatbar_tca | 0.000037 | HC=1, LR=1, NR=1 | 0.000037 | 0.000037 | 0.000037 |
| soma | sk | gskbar_sk | 0.001 | HC=1, LR=1.03, NR= 0.7878 | **0.001** | **0.00103** | **≈ 0.0007878** |
| soma | bk | gkbar_bk | 0.0006 | HC=1, LR=1, NR=1 | 0.0006 | 0.0006 | 0.0006 |
| **gcldend** | ichan2 | gnatbar_ichan2 | 0.018 | HC=1, LR=1.03, NR= 0.7878 | **0.018** | **0.01854** | **≈ 0.014182** |
| gcldend | ichan2 | gkfbar_ichan2 | 0.004 | HC=1, LR=1.15, NR=1.3 | **0.004** | **0.0046** | **0.0052** |
| gcldend | ichan2 | gksbar_ichan2 | 0.006 | HC=1, LR=1.15, NR=1.3 | **0.006** | **0.0069** | **0.0078** |
| gcldend | ichan2 | gl_ichan2 | 2.27E-05 | HC=1, LR=1, NR=1 | 2.27E-05 | 2.27E-05 | 2.27E-05 |
| gcldend | nca | gncabar_nca | 0.003 | HC=1, LR=1, NR=1 | 0.003 | 0.003 | 0.003 |
| gcldend | lca | glcabar_lca | 0.0075 | HC=1, LR=1, NR=1 | 0.0075 | 0.0075 | 0.0075 |
| gcldend | tca | gcatbar_tca | 0.000075 | HC=1, LR=1, NR=1 | 0.000075 | 0.000075 | 0.000075 |
| gcldend | sk | gskbar_sk | 0.0004 | HC=1, LR=1, NR=1 | **0.0004** | **0.0004** | **0.0004** |
| gcldend | bk | gkbar_bk | 0.0006 | HC=1, LR=1, NR=1 | 0.0006 | 0.0006 | 0.0006 |
| **pdend** | ichan2 | gnatbar_ichan2 | 0.013 | HC=1, LR=1.03, NR= 0.7878 | **0.013** | **0.01339** | **≈ 0.010242** |
| pdend | ichan2 | gkfbar_ichan2 | 0.004 | HC=1, LR=1.15, NR=1.3 | **0.004** | **0.0046** | **0.0052** |
| pdend | ichan2 | gksbar_ichan2 | 0.006 | HC=1, LR=1.15, NR=1.3 | **0.006** | **0.0069** | **0.0078** |
| pdend | ichan2 | gl_ichan2 | 2.27E-05 | HC=1, LR=1, NR=1 | 2.27E-05 | 2.27E-05 | 2.27E-05 |
| pdend | nca | gncabar_nca | 0.001 | HC=1, LR=1, NR=1 | 0.001 | 0.001 | 0.001 |
| pdend | lca | glcabar_lca | 0.0075 | HC=1, LR=1, NR=1 | 0.0075 | 0.0075 | 0.0075 |
| pdend | tca | gcatbar_tca | 0.00025 | HC=1, LR=1, NR=1 | 0.00025 | 0.00025 | 0.00025 |
| pdend | sk | gskbar_sk | 0.0002 | HC=1, LR=1, NR=1 | **0.0002** | **0.0002** | **0.0002** |
| pdend | bk | gkbar_bk | 0.001 | HC=1, LR=1, NR=1 | 0.001 | 0.001 | 0.001 |
| **mdend** | ichan2 | gnatbar_ichan2 | 0.008 | HC=1, LR=1.03, NR= 0.7878 | **0.008** | **0.0082** | **≈ 0.0063** |
| mdend | ichan2 | gkfbar_ichan2 | 0.001 | HC=1, LR=1.15, NR=1.3 | **0.001** | **0.00115** | **0.0013** |
| mdend | ichan2 | gksbar_ichan2 | 0.006 | HC=1, LR=1.15, NR=1.3 | **0.006** | **0.0069** | **0.0078** |
| mdend | ichan2 | gl_ichan2 | 2.27E-05 | HC=1, LR=1, NR=1 | 2.27E-05 | 2.27E-05 | 2.27E-05 |
| mdend | nca | gncabar_nca | 0.001 | HC=1, LR=1, NR=1 | 0.001 | 0.001 | 0.001 |
| mdend | lca | glcabar_lca | 0.0005 | HC=1, LR=1, NR=1 | 0.0005 | 0.0005 | 0.0005 |
| mdend | tca | gcatbar_tca | 0.0005 | HC=1, LR=1, NR=1 | 0.0005 | 0.0005 | 0.0005 |
| mdend | sk | gskbar_sk | 0 | HC=1, LR=1, NR=1 | **0** | **0** | **0** |
| mdend | bk | gkbar_bk | 0.0024 | HC=1, LR=1, NR=1 | 0.0024 | 0.0024 | 0.0024 |
| **ddend** | ichan2 | gnatbar_ichan2 | 0 | HC=1, LR=1.03, NR= 0.7878 | **0** | **0** | **0** |
| ddend | ichan2 | gkfbar_ichan2 | 0.001 | HC=1, LR=1.15, NR=1.3 | **0.001** | **0.00115** | **0.0013** |
| ddend | ichan2 | gksbar_ichan2 | 0.008 | HC=1, LR=1.15, NR=1.3 | 0.008 | **0.0092** | **0.0104** |
| ddend | ichan2 | gl_ichan2 | 2.27E-05 | HC=1, LR=1, NR=1 | 2.27E-05 | 2.27E-05 | 2.27E-05 |
| ddend | nca | gncabar_nca | 0.001 | HC=1, LR=1, NR=1 | 0.001 | 0.001 | 0.001 |
| ddend | lca | glcabar_lca | 0 | HC=1, LR=1, NR=1 | 0 | 0 | 0 |
| ddend | tca | gcatbar_tca | 0.001 | HC=1, LR=1, NR=1 | 0.001 | 0.001 | 0.001 |
| ddend | sk | gskbar_sk | 0 | HC=1, LR=1.03, NR= 0.7878 | **0** | **0** | **0** |
| ddend | bk | gkbar_bk | 0.0024 | HC=1, LR=1, NR=1 | 0.0024 | 0.0024 | 0.0024 |

Supplementary Table 5: Basket Cell Conductance Parameters within HC, LR, and NR

| **Section** | **Mechanism** | **Parameter** | **Scaling Factor** | **HC** | **LR** | **NR** |
| --- | --- | --- | --- | --- | --- | --- |
| All | ka | gkabar_ka | HC=1, LR=0.8, NR=0.9 | 0.00015 × 1.0 | 0.00015 × 0.8 | 0.00015 × 0.9 |
|  | nca | gncabar_nca | HC=1, LR=1, NR=1 | 0.0008 | 0.0008 | 0.0008 |
|  | lca | glcabar_lca | HC=1, LR=1, NR=1 | 0.005 | 0.005 | 0.005 |
|  | sk | gskbar_sk | HC=1, LR=1, NR=1 | 0.000002 | 0.000002 | 0.000002 |
|  | bk | gkbar_bk | HC=1, LR=1, NR=1 | 0.0002 | 0.0002 | 0.0002 |
| Soma | ichan2 | gnatbar_ichan2 | HC=1, LR=1.8, NR=0.2 | 0.12 × 1.0 | 0.12 × 1.8 | 0.12 × 0.2 |
|  |  | gkfbar_ichan2 | HC=1, LR=0.8, NR=0.85 | 0.013 × 1.0 | 0.013 × 0.8 | 0.013 × 0.85 |
|  |  | gl_ichan2 | HC=1, LR=1, NR=1 | 0.00018 | 0.00018 | 0.00018 |
| adend | ichan2 | gnatbar_ichan2 | HC=1, LR=1.8, NR=0.2 | 0.12 × 1.0 | 0.12 × 1.8 | 0.12 × 0.2 |
|  |  | gkfbar_ichan2 | HC=1, LR=0.8, NR=0.85 | 0.013 × 1.0 | 0.013 × 0.8 | 0.013 × 0.85 |
|  |  | gl_ichan2 | HC=1, LR=1, NR=1 | 0.00018 | 0.00018 | 0.00018 |
| bdend | ichan2 | gnatbar_ichan2 | HC=1, LR=1.8, NR=0.2 | 0.0 × 1.0 | 0.0 × 1.8 | 0.0 × 0.2 |
|  |  | gkfbar_ichan2 | HC=1, LR=0.8, NR=0.85 | 0.0 × 1.0 | 0.0 × 0.8 | 0.0 × 0.85 |
|  |  | gl_ichan2 | HC=1, LR=1, NR=1 | 0.00018 | 0.00018 | 0.00018 |
| cdend | ichan2 | gnatbar_ichan2 | HC=1, LR=1.8, NR=0.2 | 0.0 × 1.0 | 0.0 × 1.8 | 0.0 × 0.2 |
|  |  | gkfbar_ichan2 | HC=1, LR=0.8, NR=0.85 | 0.0 × 1.0 | 0.0 × 0.8 | 0.0 × 0.85 |
|  |  | gl_ichan2 | HC=1, LR=1, NR=1 | 0.00018 | 0.00018 | 0.00018 |
| ddend | ichan2 | gnatbar_ichan2 | HC=1, LR=1.8, NR=0.2 | 0.0 × 1.0 | 0.0 × 1.8 | 0.0 × 0.2 |
|  |  | gkfbar_ichan2 | HC=1, LR=0.8, NR=0.85 | 0.0 × 1.0 | 0.0 × 0.8 | 0.0 × 0.85 |
|  |  | gl_ichan2 | HC=1, LR=1, NR=1 | 0.00018 | 0.00018 | 0.00018 |

Supplementary Table 6: Mossy Cell Conductance Parameters within HC, LR, and NR

| **Location** | **Channel** | **Parameter Name** | **Value for HC, LR and NR** |
| --- | --- | --- | --- |
| **All sections** | ccanl | catau_ccanl | 10 |
|  |  | caiinf_ccanl | 5.00E-06 |
|  | ka | gkabar_ka | 1.00E-05 |
|  | nca | gncabar_nca | 8.00E-05 |
|  | lca | glcabar_lca | 6.00E-04 |
|  | sk | gskbar_sk | 0.016 |
|  | bk | gkbar_bk | 0.0165 |
|  | ih | ghyfbar_ih | 5.00E-06 |
|  |  | ghysbar_ih | 5.00E-06 |
| **Soma** | ichan2 | gnatbar_ichan2 | 0.12 |
|  |  | gkfbar_ichan2 | 0.0005 |
|  |  | gl_ichan2 | 1.10E-05 |
|  |  | cm | 0.6 |
| **Proximal dendrites (pdend)** | ichan2 | gnatbar_ichan2 | 0.12 |
|  |  | gkfbar_ichan2 | 0.0005 |
|  |  | gl_ichan2 | 4.40E-05 |
|  |  | cm | 2.4 |
| **Distal dendrites (ddend)** | ichan2 | gnatbar_ichan2 | 0 |
|  |  | gkfbar_ichan2 | 0 |
|  |  | gl_ichan2 | 4.40E-05 |
|  |  | cm | 2.4 |

Supplementary Table 7: Mossy Cell Conductance Parameters within HC, LR, and NR

| **Ion Channel** | **Parameter** | **Value** | **Applied to** |
| --- | --- | --- | --- |
| ccanl | catau_ccanl | 10 | all compartments |
|  | caiinf_ccanl | 5.00E-06 |  |
| ka | gkabar_ka | 0.0008 | all compartments |
| nca | gncabar_nca | 0 | all compartments |
| lca | glcabar_lca | 0.0015 | all compartments |
| sk | gskbar_sk | 0.003 | all compartments |
| bk | gkbar_bk | 0.003 | all compartments |
| ih | ghyfbar_ih | 1.50E-05 | all compartments |
|  | ghysbar_ih | 1.50E-05 |  |
| ichan2 (Na, Kf) | gnatbar_ichan2 | 0.2 | soma, proximal dendrites |
|  | gkfbar_ichan2 | 0.006 |  |
|  | gl_ichan2 | 3.60E-05 | all compartments |
|  | cm | 1.1 | all compartments |

Supplementary Table 8 : Synaptic Parameters for Granule Cell (GC), Basket Cell (BC), Mossy Cell (MC), and HIPP Interneurons.

| **Cell Type** | **Compartment** | **Synapse Type** | **Receptor** | **Tau1 (ms)** | **Tau2 (ms)** | **Reversal Potential (mV)** |
| --- | --- | --- | --- | --- | --- | --- |
| BasketCell | bcdend1 | PP | AMPA | 2 | 6.3 | 0 |
| BasketCell | bcdend2 | PP | AMPA | 2 | 6.3 | 0 |
| BasketCell | bcdend1 | GC | AMPA | 0.3 | 0.6 | 0 |
| BasketCell | bcdend2 | GC | AMPA | 0.3 | 0.6 | 0 |
| BasketCell | bcdend3 | GC | AMPA | 0.3 | 0.6 | 0 |
| BasketCell | bcdend4 | GC | AMPA | 0.3 | 0.6 | 0 |
| BasketCell | bcdend1 | MC | AMPA | 0.9 | 3.6 | 0 |
| BasketCell | bcdend2 | MC | AMPA | 0.9 | 3.6 | 0 |
| BasketCell | bcdend1 | BC | GABA | 0.16 | 1.8 | -70 |
| BasketCell | bcdend2 | BC | GABA | 0.16 | 1.8 | -70 |
| BasketCell | bcdend1 | HIPP | GABA | 0.4 | 5.8 | -70 |
| BasketCell | bcdend2 | HIPP | GABA | 0.4 | 5.8 | -70 |
| MossyCell | mcdend1 | PP | AMPA | 1.5 | 5.5 | 0 |
| MossyCell | mcdend2 | PP | AMPA | 1.5 | 5.5 | 0 |
| MossyCell | mcdend3 | PP | AMPA | 1.5 | 5.5 | 0 |
| MossyCell | mcdend4 | PP | AMPA | 1.5 | 5.5 | 0 |
| MossyCell | mcdend1 | GC | AMPA | 0.5 | 6.2 | 0 |
| MossyCell | mcdend2 | GC | AMPA | 0.5 | 6.2 | 0 |
| MossyCell | mcdend3 | GC | AMPA | 0.5 | 6.2 | 0 |
| MossyCell | mcdend4 | GC | AMPA | 0.5 | 6.2 | 0 |
| MossyCell | mcdend1 | MC | AMPA | 0.45 | 2.2 | 0 |
| MossyCell | mcdend2 | MC | AMPA | 0.45 | 2.2 | 0 |
| MossyCell | mcdend3 | MC | AMPA | 0.45 | 2.2 | 0 |
| MossyCell | mcdend4 | MC | AMPA | 0.45 | 2.2 | 0 |
| MossyCell | soma | BC | GABA | 0.3 | 3.3 | -70 |
| MossyCell | mcdend1 | HIPP | GABA | 0.5 | 6 | -70 |
| MossyCell | mcdend2 | HIPP | GABA | 0.5 | 6 | -70 |
| MossyCell | mcdend3 | HIPP | GABA | 0.5 | 6 | -70 |
| MossyCell | mcdend4 | HIPP | GABA | 0.5 | 6 | -70 |
| HIPP | hcdend1 | GC | AMPA | 0.3 | 0.6 | 0 |
| HIPP | hcdend2 | GC | AMPA | 0.3 | 0.6 | 0 |
| HIPP | hcdend3 | GC | AMPA | 0.3 | 0.6 | 0 |
| HIPP | hcdend4 | GC | AMPA | 0.3 | 0.6 | 0 |
| HIPP | hcdend1 | MC | AMPA | 0.9 | 3.6 | 0 |
| HIPP | hcdend2 | MC | AMPA | 0.9 | 3.6 | 0 |
| HIPP | hcdend3 | MC | AMPA | 0.9 | 3.6 | 0 |
| HIPP | hcdend4 | MC | AMPA | 0.9 | 3.6 | 0 |
